# BIG-TB: A benchmark for prediction and interpretability of sequence-based machine learning using *Mycobacterium tuberculosis* genomes

**DOI:** 10.64898/2026.01.30.702134

**Authors:** Mahbuba Tasmin, Mohanty Saishradha, Sanjana Kulkarni, Maha R. Farhat, Anna G. Green

**Affiliations:** Manning College of Information and Computer Sciences, University of Massachusetts, 140 Governors Drive, Amherst, MA 01002; Department of Biomedical Informatics, Harvard Medical School, 25 Shattuck St, Boston, MA 02115; Division of Pulmonary & Critical Care, Massachusetts General Hospital, 55 Fruit St, Boston, MA 02114

## Abstract

**Motivation:** Foundation models aim to learn useful representations of biological sequences, but are hindered by a lack of benchmarks to jointly evaluate predictive performance and variant attribution on diverse tasks. To this end, we present the Benchmark for Interpretable prediction from Genomes of Tuberculosis, BIG-TB.

**Results:** We curate over 17,000 genomes with high-quality sequencing data and experimentally measured antibiotic resistance phenotypes, combined with a curated list of WHO-catalogue of resistance-associated variants, and provide these data in an ML-ready format. BIG-TB defines two tasks: (1) predictive performance of antibiotic resistance phenotypes, and (2) attribution of predictions to known resistance-conferring variants. Using our tasks, we evaluate DNA- and protein-based foundation models, showing that DNA-based foundation models do not yet outperform simple machine learning baselines (mean test AUC = 0.888 vs. 0.865 for best CNN variant vs. best Evo2 variant across drugs). In a lineage-informed evaluation across DNA and protein modalities, predictive signal for well-characterized drugs was largely preserved (mean AUC 0.846–0.963). Across both evaluation settings, models with the highest predictive performance did not necessarily achieve the strongest recovery of resistance variants, underscoring the value of evaluating both predictive accuracy and biological interpretability. Overall, BIG-TB provides a reusable resource for developing sequence representations that are both accurate and mechanistically meaningful.

**Availability:** Code is available at SAGE-Lab-UMass/Big-TB-benchmark; data are available at figshare.

**Supplementary information:** Supplementary data are available at *Bioinformatics* online.

## 1 Introduction

Foundation models for biological sequences promise broadly reusable representations of protein [1–3] and DNA [4–8] sequences. Increasing numbers of benchmarks have been introduced to evaluate the accuracy of these models across ranges of tasks and organisms [4, 6–10]. Recent evaluations demonstrate that pre-trained sequence encoders can improve predictive accuracy across diverse biological applications, including regulatory genomics, variant-effect prediction, and protein function modeling, [5, 11, 12] though other studies find no or marginal utility of foundation models compared to traditional models [13, 14].

But, predictive accuracy alone is not enough; interpretability, which aims to understand which signals the model uses to make predictions [15–17], is a central goal in biological sequence models [18]. Interpretability methods can be used to discover and prioritize sequence features underlying biological processes of interest, such as gene regulation [19–21], gene–gene and gene–environment interactions [22], and antibiotic resistance [23, 24]. However, it is difficult to evaluate interpretability methods because the ground-truth causal sequence features are often not known in advance [18].

Tuberculosis, the disease caused by *Mycobacterium tuberculosis*, remains a highly burdensome infectious disease [25]; rapid, interpretable predictions from sequence can inform treatment and surveillance [26, 27]. We provide a benchmark dataset to simultaneously evaluate sequence models for predictive performance and agreement between model attributions and curated resistance-associated variants. Our phenotype of interest is antibiotic resistance because, in *M. tuberculosis*, extensive genotype–phenotype studies have produced both isolate-level drug-susceptibility data and curated variant-level evidence on resistance association [28–31]. Existing tools for predicting *M. tuberculosis* drug resistance from genome sequence, including catalogue-based systems such as Mykrobe, GenTB, and TBProfiler, are designed primarily to generate resistance calls for input isolates in clinical or surveillance workflows [32–34]. While these tools provide valuable genotype-to-phenotype predictions, they are not designed as standardized evaluation frameworks for systematically comparing learned sequence representations and their biological interpretability.

BIG-TB addresses this need by providing a unified benchmark in which researchers can evaluate not only whether a sequence-based model accurately predicts antibiotic resistance, but also whether its predictions rely on genomic positions associated with known resistance mechanisms. By combining curated genotype–phenotype data with WHO-catalogued resistance variants, BIG-TB enables quantitative comparison of predictive performance and attribution-based variant recovery across model architectures. Furthermore, lineage-holdout evaluation allows researchers to examine whether these properties are maintained across distinct M. tuberculosis genetic backgrounds. These complementary evaluations provide a framework for investigating the relationship between predictive accuracy, biological interpretability, and robustness to population structure.

Here, we introduce the Benchmark for Interpretable prediction from Genomes of Tuberculosis, BIG-TB, a publicly available dataset and benchmark that unifies prediction and interpretability evaluation for antibiotic resistance in *M. tuberculosis* (Figure 1). Specifically, BIG-TB (1) provides curated genotype and phenotype labels for over 17,000 clinical isolates, (2) links genotype data to WHO-catalogued resistance-associated variants, (3) defines a WHO-catalogued resistance variant recovery task for quantitative interpretability (building on the framework introduced in 35) and (4) standardizes phenotype prediction and attribution-based variant recovery within the same dataset. Using Big-TB, we systematically compare classical machine learning, convolutional and transformer architectures, and pretrained sequence representations from Evo2 and ESM-2. We further investigate how alternative embedding representations and dimensionality reduction affect downstream performance. To assess robustness beyond random train–test partitioning, we additionally use lineage-holdout splits, which test prediction and WHO-catalogued resistance variant recovery in held-out *M. tuberculosis* genetic backgrounds. This benchmark provides a unified, publicly reusable resource to guide the development of accurate and interpretable sequence-based models for biological prediction.

**Figure 1:**
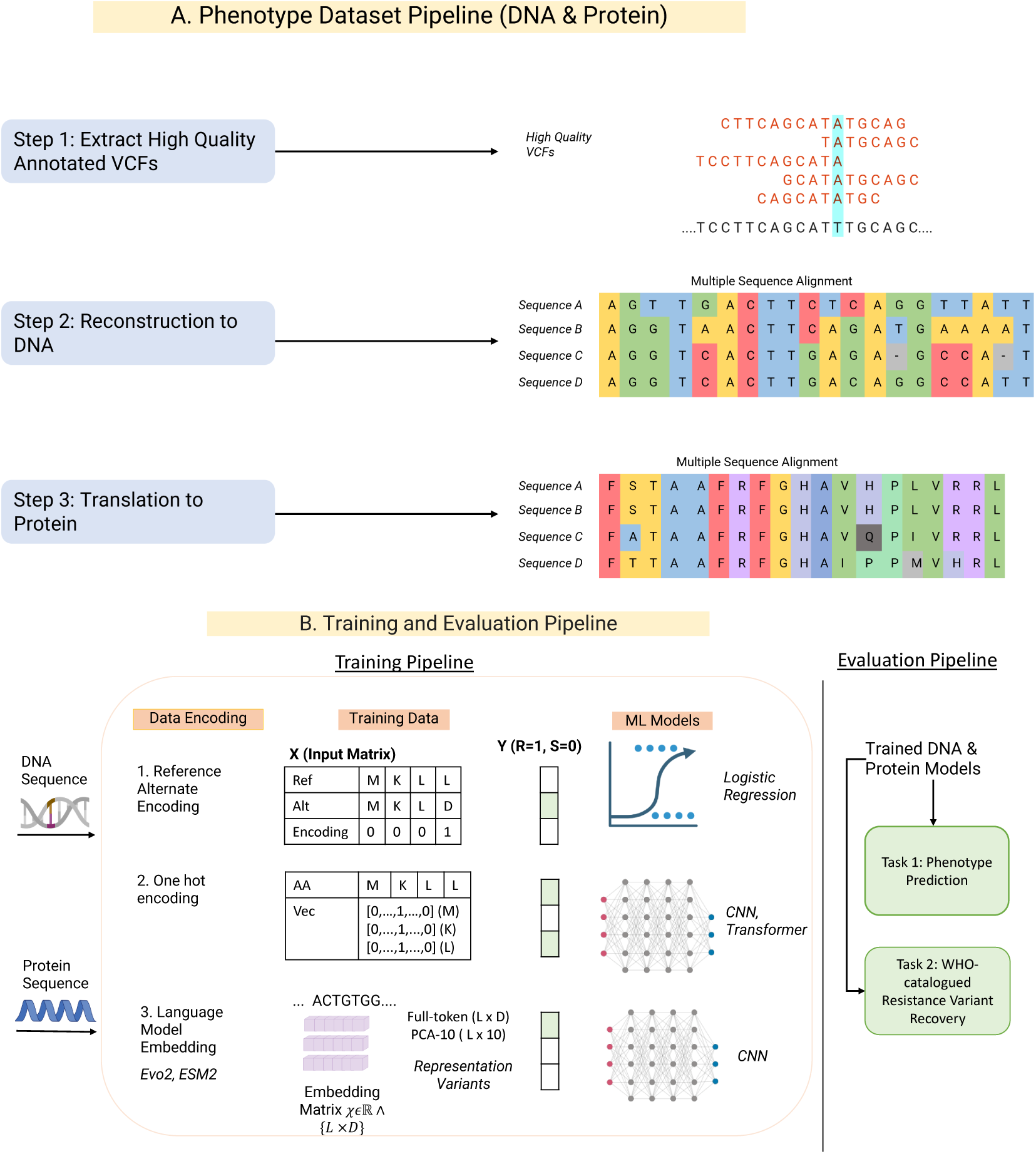
BIG-TB genotype and evaluation pipeline. (A) Genotype dataset pipeline: High-quality VCFs are reconstructed into gene-centric DNA sequences, aligned into multiple sequence alignments (MSAs), and translated into protein MSAs. (B) Training and evaluation pipeline: DNA and protein sequences are encoded using three representation strategies—(1) reference–alternate encoding for mutation indicators used in logistic regression, (2) one-hot encoding for CNN and Transformer models, and (3) contextual embeddings from pretrained language models (Evo2 for DNA, ESM2 for protein) processed via CNNs. All model families are trained to predict resistance phenotypes, and their position-level attributions are evaluated for recovery of WHO-catalogued resistance variants.

## 2 Results

### 2.1 Dataset Overview

We compiled resistance phenotypes and genomic variants for 17,942 clinical *M. tuberculosis* isolates (Table 1). For rifampicin, 94.7% of phenotypically resistant isolates carry at least one WHO-catalogued resistance variant, but for other drugs this fraction ranges from 67.5% (moxifloxacin) to 90.8% (levofloxacin). Catalogue lookup alone would miss a substantial share of resistance, justifying the need for ML models, rather than rule-based methods to predict resistance.

**Table 1:** Summary of phenotype and mutation variant statistics per drug. R/S: phenotyped resistant and susceptible isolates (isolates without labels excluded). WHO variant columns give catalogued positions per drug, split into resistance-associated (groups 1–2) and uncertain or not associated (groups 3–5; Table 6). R Isolates w/ WHO Variant: fraction of resistant isolates carrying at least one group 1–2 variant. WHO R Variants Observed: fraction of group 1–2 positions observed at least once in the data.

| Drug | Phenotyped Isolates (#) |  |  | WHO Catalogue Variants |  |  | R Isolates w. WHO Variant (Ratio) | WHO R Variants Observed (Ratio) |
|---|---|---|---|---|---|---|---|---|
|  | Resistant | Susceptible | Resistance Rate | Total | Assoc. w. Resistance | Not/Uncertain |  |  |
| Isoniazid | 5825 | 11610 | 0.334 | 7269 | 142 | 7127 | 0.906 | 0.316 |
| Rifampicin | 4869 | 12712 | 0.277 | 7266 | 136 | 7130 | 0.947 | 0.58 |
| Ethambutol | 2945 | 12165 | 0.195 | 7109 | 13 | 7096 | 0.792 | 1.00 |
| Pyrazinamide | 2131 | 10772 | 0.165 | 2562 | 341 | 2221 | 0.803 | 0.75 |
| Streptomycin | 2416 | 5266 | 0.315 | 3052 | 159 | 2893 | 0.732 | 0.635 |
| Amikacin | 707 | 2998 | 0.191 | 2183 | 4 | 2179 | 0.696 | 1.00 |
| Capreomycin | 675 | 3179 | 0.175 | 2648 | 70 | 2578 | 0.692 | 0.214 |
| Kanamycin | 703 | 2873 | 0.197 | 2233 | 8 | 2225 | 0.821 | 1.00 |
| Moxifloxacin | 388 | 2480 | 0.135 | 2717 | 18 | 2699 | 0.675 | 1.00 |
| Levofloxacin | 76 | 193 | 0.282 | 3040 | 18 | 3022 | 0.908 | 1.00 |
| Ethionamide | 690 | 1463 | 0.320 | 2739 | 286 | 2453 | 0.787 | 0.49 |

### 2.2 Benchmarking foundation models

We ask whether foundation models can perform accurate antibiotic resistance phenotype prediction and can recover known WHO-catalogued resistance variants. To this end, we evaluate single–drug and multi-drug models on two standardized tasks: Task 1 (Phenotype prediction), and Task 2 (WHO-catalogued resistance variant recovery). We compare to logistic regression as a simple machine learning baseline, and against published CNN models on one-hot encoded sequences as a deep learning baseline [36].

### 2.3 Task 1: Phenotype Prediction

We compare both protein and DNA models against their respective Logistic Regression baseline [36] using paired Wilcoxon signed-rank tests for eleven drugs (Table 15) across identical cross-validation folds (Figure 2). For DNA models, we evaluate both single-drug (SD) models, which predict one phenotype per model, and multi-drug (MD) models, which seek to predict all resistance phenotypes simultaneously via multi-task learning [36]. For evaluation across lineage-aware data splits, see section 2.6. The Evo2 models described below use a frozen encoder plus trained CNN layers. Due to computational restrictions as described in appendix G.3 we perform fine-tuning of Evo2 only for lineage-aware splits. Evaluations are performed on isolate-level cross-validation or held-out test splits as specified below (Section 4.4); per-drug train/test counts are given in Table 14.

**Figure 2:**
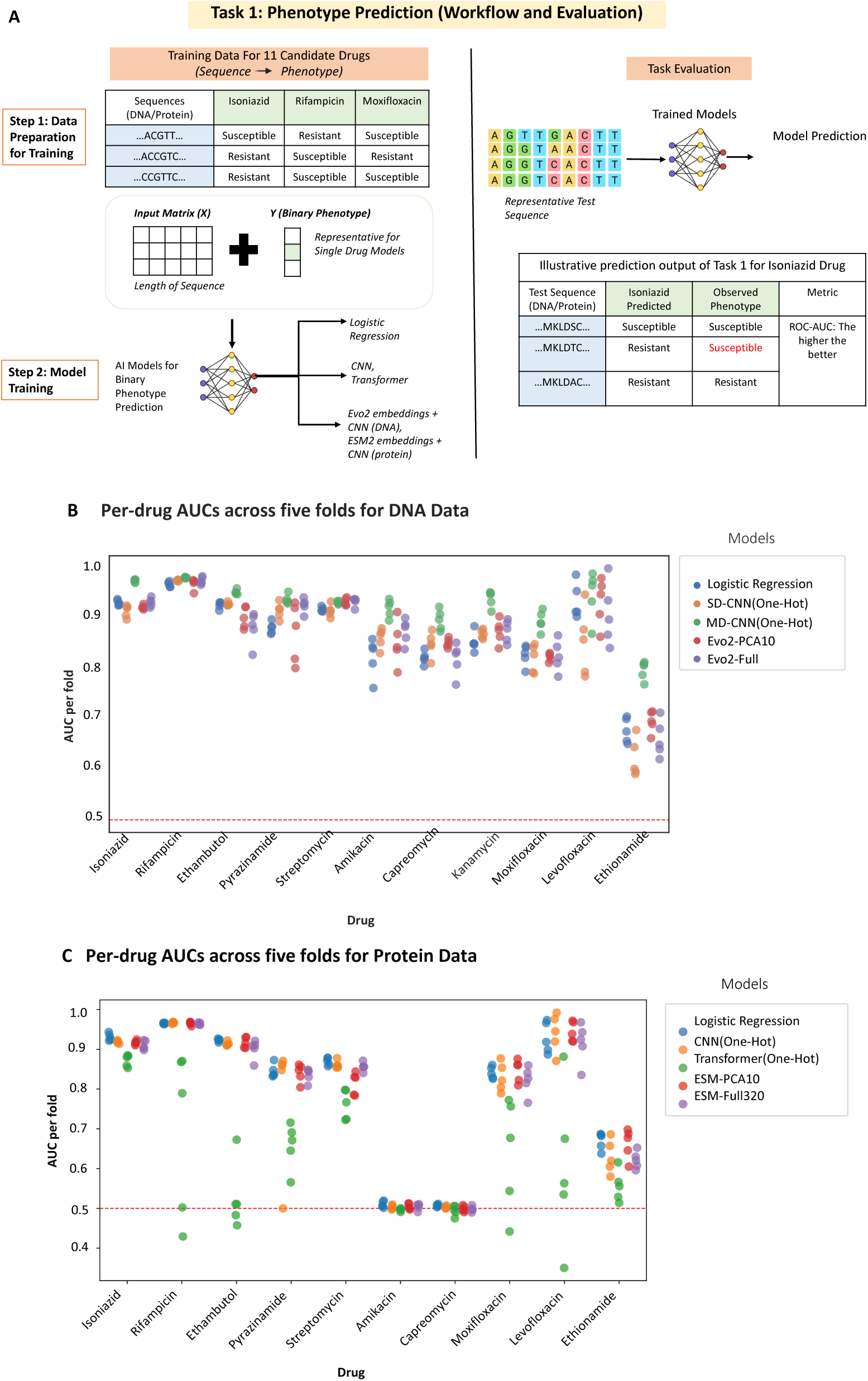
Overview of Task 1: phenotype prediction from DNA and protein sequence features. (A) Workflow showing model training and evaluation setup. (B) DNA-based model performance in cross-validation across 11 drugs; each point represents ROC–AUC on one held-out fold per model. (C) Protein-based model performance in cross-validation across 10 drugs. Red dashed line indicates random-chance performance (AUC = 0.5).

#### Cross-validation performance

For single-drug tasks, the SD-CNN performed comparably to the baseline SD-LogReg (mean cross-validation AUC = 0.864 vs. 0.867; *p* = 0.966, Holm-corrected *p* = 1.000; Table 15), consistent with prior findings. We then benchmark genomic foundation models: SD-Evo2-CNN models trained on token-level embeddings also performed comparably to the baseline across drugs, with a mean AUC of 0.871 for SD-Evo2-CNN (*p* = 0.765) and 0.873 for SD-Evo2-CNN-PCA10 (*p* = 0.413; Holm-corrected *p* = 1.000 for both). Overall, none of the single-drug models differed significantly from SD-LogReg, and the evaluated frozen SD-Evo2-CNNs did not significantly surpass the baselines. The multi-drug MD-CNN achieved the highest mean cross-validation AUC (0.922) and was the only model to significantly outperform SD-LogReg (*p* = 0.001, Holm-corrected *p* = 0.004; Table 15).

For protein-based tasks, Logistic Regression achieved the highest mean cross-validation AUC (0.799; Figure 2), and no protein architecture significantly outperformed it (Table 16). The CNN (0.785; *p* = 0.037, Holm-corrected *p* = 0.074) and ESM-PCA10 (0.791; *p* = 0.131, Holm-corrected *p* = 0.131) performed comparably to the baseline, whereas ESM-320 (0.783) and the Transformer (0.630) performed significantly worse (Holm-corrected *p* = 0.008 for both). Amikacin and capreomycin yielded near-random AUCs (*≈* 0.5) because their main resistance determinants reside in ribosomal RNA, so the input proteins contain insufficient signal (Figure 2).

#### Test set accuracy

We then evaluate our DNA and protein models on the held-out test set (Table 2). Consistently, MD-CNN achieves the highest AUC values for the majority of drugs, with AUC *>* 0.9 for ethambutol, isoniazid, pyrazinamide, rifampicin, and streptomycin. This indicates that concatenated, multi-gene representations are particularly effective when modeling resistance mechanisms jointly across drugs, though we note that this may be attributable to signal from non-causal mutations [36, 37]. In contrast, the single-drug frozen Evo2 embeddings, using either the full representation or its PCA-compressed form, do not consistently outperform the classical one-hot encoded CNN; SD-Evo2-CNN achieved the highest test AUC only for levofloxacin (0.988), the drug with the least training data.

**Table 2:** Test AUC across first- and second-line antibiotics using nucleotide (DNA) and protein sequence inputs. DNA results (top) are based on single- or multi-drug modeling; protein results (bottom) use per-drug inputs of single genes or concatenations per known mechanisms. Protein models which exclude the major resistance-determining regions of the small and large ribosomal RNA are marked with *. Protein results for Kanamycin were not available since its locus is not protein-coding (marked with “—”).

| DNA |  |  |  |  |  |
|---|---|---|---|---|---|
| Drug | SD-LogReg (Ref–Alt) | SD-CNN (OHE) | MD-CNN (OHE) | SD-Evo2-CNN (frozen) | SD-Evo2-CNN-PCA10 (frozen) |
| Isoniazid | 0.920 | 0.917 | **0.951** | 0.933 | 0.934 |
| Rifampicin | 0.975 | 0.977 | **0.981** | 0.969 | 0.964 |
| Ethambutol | 0.915 | 0.931 | **0.936** | 0.887 | 0.835 |
| Pyrazinamide | 0.896 | **0.930** | 0.908 | 0.922 | 0.922 |
| Streptomycin | 0.924 | 0.911 | **0.938** | 0.933 | 0.927 |
| Amikacin | 0.807 | 0.845 | **0.899** | 0.863 | 0.825 |
| Capreomycin | 0.846 | **0.873** | 0.829 | 0.846 | 0.823 |
| Kanamycin | 0.834 | 0.849 | **0.875** | 0.868 | 0.863 |
| Moxifloxacin | 0.864 | 0.886 | **0.887** | 0.842 | 0.825 |
| Levofloxacin | 0.835 | 0.839 | 0.850 | **0.988** | 0.957 |
| Ethionamide | 0.708 | 0.670 | **0.709** | 0.637 | 0.708 |
| Protein |  |  |  |  |  |
| Drug | LogReg (Ref–Alt) | CNN (OHE) | Transformer (OHE) | ESM-320 | ESM-PCA10 |
| Isoniazid | **0.932** | 0.919 | 0.887 | 0.912 | 0.907 |
| Rifampicin | 0.961 | 0.961 | 0.533 | **0.969** | 0.969 |
| Ethambutol | 0.921 | **0.923** | 0.640 | 0.750 | 0.911 |
| Pyrazinamide | **0.867** | 0.820 | 0.728 | 0.851 | 0.844 |
| Streptomycin* | 0.851 | 0.840 | 0.728 | **0.858** | 0.832 |
| Amikacin* | **0.508** | 0.501 | 0.500 | 0.503 | 0.505 |
| Capreomycin* | **0.505** | 0.501 | 0.489 | 0.502 | 0.499 |
| Kanamycin* | — |  |  |  |  |
| Moxifloxacin | 0.817 | 0.807 | 0.613 | 0.804 | **0.820** |
| Levofloxacin | 0.876 | **0.997** | 0.856 | 0.864 | 0.921 |
| Ethionamide | **0.728** | 0.620 | 0.405 | 0.638 | 0.663 |

The protein sequence–based results show a more heterogeneous performance landscape across antibiotics. CNNs trained directly on one-hot encoded sequences (CNN-OHE) and models based on ESM embeddings (ESM-320 and ESM-PCA10) achieved the highest AUCs for most drugs, with CNN-OHE reaching near-perfect separation for levofloxacin (AUC = 0.997) and maintaining strong performance for ethambutol and rifampicin. Logistic Regression using Ref–Alt encodings remained competitive, particularly for isoniazid (0.932) and rifampicin (0.961). Transformers trained on one-hot inputs consistently underperformed CNNs and ESM-based models.

#### Comparison between DNA- and protein-based models

DNA-based models generally achieved higher test AUCs than protein-based models for the same drugs (Table 2), likely because they incorporate both coding and non-coding regions, including regulatory and rRNA sequences that are excluded from protein-based analyses. Protein models performed well only for drugs whose known resistance-conferring mutations occur within protein-coding genes (*rpoB*, *katG*, *embB*).

### 2.4 Task 2: WHO-catalogued resistance variant recovery

We next evaluated whether SHAP attributions recover known resistance-associated variants from the WHO 2023 mutation catalogue (Figure 3, Table 3). For each model, we measure the precision and recall of the top-10 variants ranked by SHAP analysis. Because target-set size varies substantially across drugs, we interpret Recall@10 jointly with Precision@10, and compare the values across models for the same drug only.

**Figure 3:**
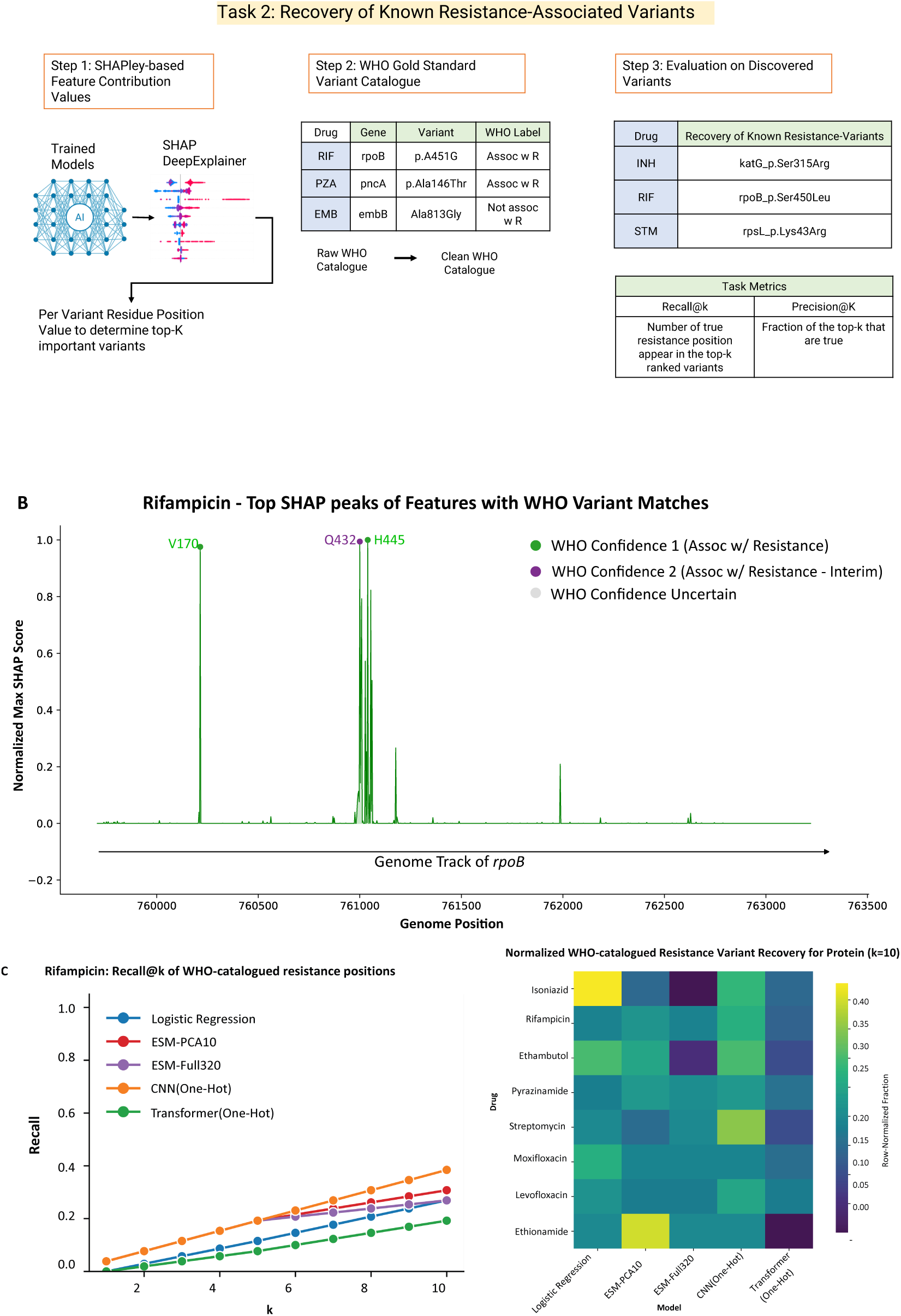
WHO-catalogued resistance variant recovery. (a) Workflow for ranking model-attributed sites against resistance-associated variants in the WHO 2023 catalogue using top-*k* evaluation. (b) Genome tracks for rifampicin showing SHAP peaks at the *rpoB* locus alongside WHO-catalogued resistance variants. (c) Left: Recall@*k* for rifampicin, the fraction of WHO target positions recovered among each model’s top-*k* SHAP-ranked sites (*k* = 1–10). Right: Recall@10 per drug, row-normalized so that each cell is a model’s Recall@10 divided by the sum across models for that drug; colours therefore show each model’s relative share of recovery within a drug. Amikacin, capreomycin, and kanamycin are not shown.

**Table 3:** Precision@10 and Recall@10 for recovery of WHO-catalogued resistance target positions across DNA-and protein-based models. Bold values indicate the best value per drug within each modality. Precision@10 and Recall@10 were computed using the best-performing cross-validation model selected in Task 1 for each drug and model. The reported values are single values obtained from the selected model and are not averaged across cross-validation folds.

| DNA |  |  |  |  |  |  |  |  |  |  |  |
| Drug | Residues recovered | SD-LogReg (Ref–Alt) |  | SD-CNN (OHE) |  | MD-CNN (OHE) |  | SD-Evo2-CNN (frozen) |  | SD-Evo2-CNN-PCA10 (frozen) |  |
|  |  | **Prec@10** | **Recall@10** | **Prec@10** | **Recall@10** | **Prec@10** | **Recall@10** | **Prec@10** | **Recall@10** | **Prec@10** | **Recall@10** |
|---|---|---|---|---|---|---|---|---|---|---|---|
| Isoniazid | 42 | **0.30** | **0.071** | **0.30** | **0.071** | 0.20 | 0.048 | 0.20 | 0.048 | 0.20 | 0.048 |
| Rifampicin | 47 | 0.30 | 0.064 | 0.50 | 0.106 | 0.30 | 0.064 | **0.40** | **0.085** | 0.00 | 0.000 |
| Ethambutol | 9 | 0.50 | 0.556 | **0.70** | **0.778** | 0.10 | 0.111 | 0.00 | 0.000 | 0.00 | 0.000 |
| Pyrazinamide | 194 | **0.80** | **0.041** | 0.50 | 0.026 | 0.00 | 0.000 | 0.60 | 0.031 | 0.60 | 0.031 |
| Streptomycin | 101 | **0.50** | **0.050** | 0.40 | 0.040 | 0.30 | 0.030 | 0.20 | 0.020 | **0.50** | **0.050** |
| Amikacin | 4 | 0.20 | 0.500 | **0.30** | **0.750** | 0.10 | 0.250 | 0.20 | 0.500 | 0.10 | 0.250 |
| Capreomycin | 16 | 0.20 | 0.125 | 0.20 | 0.125 | 0.10 | 0.063 | **0.30** | **0.188** | 0.10 | 0.063 |
| Kanamycin | 8 | **0.20** | **0.250** | 0.10 | 0.125 | 0.10 | 0.125 | **0.20** | **0.250** | **0.20** | **0.250** |
| Moxifloxacin | 14 | 0.40 | 0.286 | **0.50** | **0.357** | 0.10 | 0.071 | 0.20 | 0.143 | 0.30 | 0.214 |
| Levofloxacin | 14 | 0.40 | 0.286 | **0.50** | **0.357** | 0.10 | 0.071 | 0.10 | 0.071 | 0.20 | 0.143 |
| Ethionamide | 150 | **0.50** | **0.033** | 0.30 | 0.020 | 0.10 | 0.007 | 0.10 | 0.007 | 0.00 | 0.000 |
| Protein |  |  |  |  |  |  |  |  |  |  |  |
| Drug | Residues recovered | LogReg (Ref–Alt) |  | CNN (OHE) |  | Transformer (OHE) |  | ESM-320 (Full) |  | ESM-PCA10 |  |
|  |  | **Prec@10** | **Recall@10** | **Prec@10** | **Recall@10** | **Prec@10** | **Recall@10** | **Prec@10** | **Recall@10** | **Prec@10** | **Recall@10** |
| Isoniazid | 3 | **0.30** | **1.00** | 0.20 | 0.67 | 0.10 | 0.33 | 0.00 | 0.00 | 0.10 | 0.33 |
| Rifampicin | 26 | 0.70 | 0.27 | **1.00** | **0.38** | 0.50 | 0.19 | 0.70 | 0.27 | 0.80 | 0.31 |
| Ethambutol | 6 | **0.60** | **1.00** | **0.60** | **1.00** | 0.20 | 0.33 | 0.10 | 0.17 | 0.50 | 0.83 |
| Pyrazinamide | 95 | 0.80 | 0.08 | **1.00** | **0.11** | 0.70 | 0.07 | 0.90 | 0.09 | **1.00** | **0.11** |
| Streptomycin | 10 | 0.40 | 0.40 | **0.70** | **0.70** | 0.20 | 0.20 | 0.40 | 0.40 | 0.30 | 0.30 |
| Amikacin | 0 | 0.00 | 0.00 | 0.00 | 0.00 | 0.00 | 0.00 | 0.00 | 0.00 | 0.00 | 0.00 |
| Capreomycin | 2 | 0.00 | 0.00 | **0.10** | **0.50** | 0.00 | 0.00 | 0.00 | 0.00 | **0.10** | **0.50** |
| Kanamycin | – | – | – | – | – | – | – | – | – | – | – |
| Moxifloxacin | 10 | **0.70** | **0.70** | 0.50 | 0.50 | 0.40 | 0.40 | 0.50 | 0.50 | 0.50 | 0.50 |
| Levofloxacin | 10 | 0.60 | 0.60 | **0.70** | **0.70** | 0.50 | 0.50 | 0.50 | 0.50 | 0.50 | 0.50 |
| Ethionamide | 10 | 0.10 | 0.10 | 0.10 | 0.10 | 0.00 | 0.00 | 0.10 | 0.10 | **0.20** | **0.20** |

For DNA models (Table 19; Table 23), SD-CNN and SD-Evo2-CNN (frozen) were statistically indistinguishable from SD-LogReg, whereas MD-CNN (Precision@10/Recall@10 Δ = *−*0.255*/ −* 0.129, adjusted *p* = 0.020*/*0.020) and SD-Evo2-CNN-PCA10 (frozen) (Δ = *−*0.191*/ −* 0.110, adjusted *p* = 0.023*/*0.023) recovered significantly fewer WHO-catalogued positions. MD-CNN’s weaker recovery despite its strong Task 1 performance shows that higher prediction accuracy does not imply stronger recovery of known resistance determinants.

For protein models (Table 20; Table 23), CNN, ESM-320 and ESM-PCA10 did not differ significantly from Logistic Regression, whereas the Transformer (Precision@10/Recall@10 Δ = *−*0.160*/−*0.212, adjusted *p* = 0.047*/*0.046) recovered significantly fewer WHO-catalogued positions, consistent with its weak Task 1 performance. Recovery was strongest for ethambutol and isoniazid, where Logistic Regression recovered all WHO-catalogued targets (Recall@10 = 1.0; also 1.0 for the CNN on ethambutol), and for the fluoroquinolones (Recall@10 up to 0.70 across *gyrA* and *gyrB*).

#### Linking predictive accuracy and WHO-catalogued resistance variant recovery

Predictive accuracy did not reliably indicate recovery of WHO-catalogued resistance variants in both modalities. Among DNA models, MD-CNN had the highest predictive accuracy, yet SD-CNN agreed more closely with the drug-specific WHO reference set; for the multi-drug models, many top-ranked features were variants conferring resistance to other antibiotics rather than direct mechanistic causes of resistance to the drug in question. Likewise, among protein models, CNN-OHE model is not uniformly superior in recovering known-resistance variants among drug targets; where Logistic-Regression achieves complete recovery for isoniazid and ethambutol.

### 2.5 Ablation Study of Foundation Model Embedding Variants

We investigate how different strategies for representing and compressing token-level embeddings from Evo2 (DNA) and ESM–2 (protein) affect downstream resistance prediction performance. Specifically, we compare full per-token embeddings, PCA-*k* projections, channel-mean embeddings, and sequence-mean embeddings, which differ in their preservation of positional information and contextual features (Table 13). Detailed experimental results, including per-drug performance comparisons and ablation heatmaps, are presented in Supplementary Section I.1.

### 2.6 Robustness to lineage structure

We next evaluated robustness to *M. tuberculosis* population structure using lineage-holdout splits (Section 4.5); for DNA foundation models, this evaluation used both frozen and LoRA fine-tuned Evo-2 embeddings (Section G.3).

#### Task 1: predictive performance

In DNA evaluations, mean ΔAUC under lineage holdout ranged from *−*0.048 (SD-CNN) to *−*0.036 (SD-LogReg) across the four architectures (Table 4, Supplementary Figure 7). Only SD-Evo2-CNN (frozen) declined significantly (ΔAUC = *−*0.046, Holm-corrected *p* = 0.023), largely because of pyrazinamide (0.922 to 0.718). LoRA fine-tuning did not consistently improve Evo2 under lineage holdout and reduced AUC for several drugs. These results suggest that task-specific adaptation to the training lineages can reduce the robustness of pretrained Evo2 representations to held-out lineages.

**Table 4:** Mean test ROC-AUC across eligible lineage-holdout splits (Section 4.5) for DNA and protein inputs; “—” marks unavailable results. The **Mean** row gives each model’s lineage-split minus random-split AUC (Table 2), averaged over drugs with both values.

| DNA |  |  |  |  |  |  |
|---|---|---|---|---|---|---|
| Drug | SD-LogReg (Ref–Alt) | SD-CNN (OHE) | SD-Evo2-CNN (finetuned) | SD-Evo2-CNN (frozen) | SD-Evo2-CNN-PCA10 (frozen) | Eligible held-out lineages |
| Isoniazid | **0.916** | 0.887 | 0.797 | 0.899 | 0.878 | 1, 2, 3, 4 |
| Rifampicin | 0.961 | **0.963** | 0.962 | 0.960 | 0.937 | 1, 2, 3, 4 |
| Ethambutol | **0.918** | 0.909 | 0.476 | 0.632 | 0.884 | 1, 2, 3, 4 |
| Pyrazinamide | 0.816 | **0.846** | 0.738 | 0.718 | 0.766 | 1, 2, 3, 4 |
| Streptomycin | 0.855 | **0.871** | 0.737 | 0.838 | 0.838 | 1, 2, 3, 4 |
| Amikacin | 0.834 | **0.878** | 0.744 | 0.840 | 0.807 | 2, 4 |
| Capreomycin | 0.799 | 0.810 | 0.812 | **0.825** | **0.825** | 2, 4 |
| Kanamycin | 0.834 | 0.824 | 0.848 | 0.842 | **0.863** | 2, 4 |
| Moxifloxacin | 0.792 | 0.780 | 0.778 | **0.807** | 0.795 | 2, 4 |
| Ethionamide | 0.597 | **0.612** | 0.553 | 0.610 | 0.571 | 2, 4 |
| **Mean ΔAUC** | −0.036 | −0.048 | — | −0.046 | −0.046 |  |
| Protein |  |  |  |  |  |  |
| Drug | LogReg | CNN (OHE) | Transformer (OHE) | ESM-320 | ESM-PCA10 | Eligible held-out lineages |
| Isoniazid | **0.922** | 0.907 | 0.654 | 0.903 | 0.891 | 1, 2, 3, 4 |
| Rifampicin | 0.960 | 0.961 | 0.706 | **0.963** | 0.960 | 1, 2, 3, 4 |
| Ethambutol | **0.901** | 0.898 | 0.633 | 0.823 | 0.900 | 1, 2, 3, 4 |
| Pyrazinamide | 0.780 | **0.792** | 0.661 | 0.774 | 0.738 | 1, 2, 3, 4 |
| Streptomycin | 0.763 | **0.796** | 0.639 | 0.793 | 0.723 | 1, 2, 3, 4 |
| Amikacin | **0.507** | 0.486 | 0.497 | 0.492 | 0.494 | 2, 4 |
| Capreomycin | 0.500 | **0.501** | 0.500 | 0.483 | 0.492 | 2, 4 |
| Kanamycin |  |  | — |  |  | 2, 4 |
| Moxifloxacin | **0.808** | 0.795 | 0.607 | 0.787 | 0.798 | 2, 4 |
| Ethionamide | **0.613** | 0.584 | 0.523 | 0.505 | 0.540 | 2, 4 |
| **Mean ΔAUC** | −0.037 | −0.019 | −0.011 | −0.029 | −0.046 |  |

In protein evaluations, mean ΔAUC under lineage holdout ranged from *−*0.046 (ESM-PCA10) to *−*0.011 (Transformer) (Table 4). The declines were significant for Logistic Regression (*−*0.037), CNN (*−*0.019), and ESM-PCA10 (*−*0.046; Holm-corrected *p* = 0.020 for all three), but not for Transformer or ESM-320. Overall, these results suggest that protein-based models exhibit varying sensitivity to lineage distribution shifts. Supplementary Figure 8 shows the full per-architecture breakdown.

#### Task 2: variant recovery

We repeated Task 2 using the lineage-holdout models from section 2.6.

For DNA-based Task 2 lineage holdout, the supervised baselines showed no significant change in mean Precision@10 or Recall@10 (Table 21). Full SD-Evo2-CNN declined substantially (Δ = *−*0.118*/ −* 0.076, adjusted *p* = 0.069*/*0.031 for Precision@10/Recall@10, respectively), whereas SD-Evo2-CNN-PCA10 remained stable (Δ = *−*0.038*/*+0.014, adjusted *p* = 1.000*/*1.000). LoRA fine-tuning did not recover attribution performance: fine-tuned Evo2 reached mean Precision@10/Recall@10 of only 0.053*/*0.024, compared with 0.123*/*0.052 for frozen Evo2 (Supplementary Figure 9).

For protein-based Task 2 lineage holdout, the mean Precision@10 and Recall@10 increased for every architecture relative to the random split, but no gain was significant after correction (Table 22): LogReg (Δ = +0.028*/* + 0.028, adjusted *p* = 0.924*/*0.947), CNN (Δ = +0.016*/* + 0.033, adjusted *p* = 0.924*/*0.947), Transformer (Δ = +0.069*/* + 0.032, adjusted *p* = 0.250*/*0.947), ESM-320 (Δ = +0.097*/* + 0.166, adjusted *p* = 0.232*/*0.139), and ESM-PCA10 (Δ = +0.056*/* + 0.065, adjusted *p* = 0.250*/*0.947) (Supplementary Figure 10).

## 3 Discussion & Conclusion

We present BIG-TB, a foundation model benchmark for antibiotic resistance phenotype prediction and WHO-catalogued resistance variant recovery in *M. tuberculosis*. Our benchmark enables DNA-level representations of any genomic region, and protein-level representations of coding regions, combined with phenotype data for 11 antibiotics. We uniformly preprocessed publicly available *M. tuberculosis* sequencing data using stringent quality thresholds and linked annotated mutations to the WHO 2023 catalogue of resistance-associated variants [30]. This design enables joint evaluation of predictive accuracy (Task 1) and attribution-based recovery of curated resistance variants (Task 2) across classical machine-learning and foundation-model architectures.

Using BIG-TB, we compared the general-purpose Evo2 and ESM models with simpler machine-learning methods for antibiotic-resistance prediction. For DNA models, frozen or finetuned Evo2 representations did not consistently outperform matched single-drug models trained on one-hot or reference-alternate encodings, while the multi-drug CNN achieved the strongest performance for most drugs. For protein-based models, compact position-preserving embedding variants were competitive with simple models, but their gains were generally modest (Supplementary Figure 4a,4b). DNA-based models achieved higher predictive accuracy than protein-based models for several drugs, consistent with the contribution of non-coding and rRNA variation that is unavailable to protein-only inputs.

Expanding the DNA modeling framework to multi-drug, multi-gene prediction improved predictive accuracy while adversely affecting WHO-catalogued variant recovery. For several drugs, top-ranked features in multi-drug models were associated with resistance to other drugs or fell outside the drug-specific WHO target set, consistent with correlated resistance phenotypes in MDR/XDR isolates [36, 37]. These results caution against naïve multitask integration and motivate explicit modeling of shared or correlated resistance pathways [37]. They also urge caution when incorporating additional loci into predictive models – as any locus with correlated, non-causal variation may result in improved predictive accuracy at the expense of biological fidelity.

The limited gains from genomic foundation models such as Evo2 over simpler architectures may partly reflect the constrained nature of the prediction task. Our inputs are restricted to predefined loci already known to harbor resistance-associated variation, providing a strong prior over the relevant genomic regions. Consequently, simpler models may have sufficient capacity to capture much of the discriminative signal. Fine-tuning Evo2 also did not consistently improve over its frozen representations (Tables 3 and 5), with particularly pronounced degradation for Task 2. One possible explanation is that drug-specific fine-tuning encourages the model to exploit correlations across the concatenated resistance-associated loci that are predictive of the strain-level phenotype but are not directly related to causal resistance mechanisms. These results highlight room for improvements on the utility of genomic foundation models for resistance prediction. Downstream performance may be further improved through more targeted optimization of the representation layer, pooling strategy, fine-tuning configuration, and classifier hyperparameters for each foundation model and prediction task.

**Table 5:**
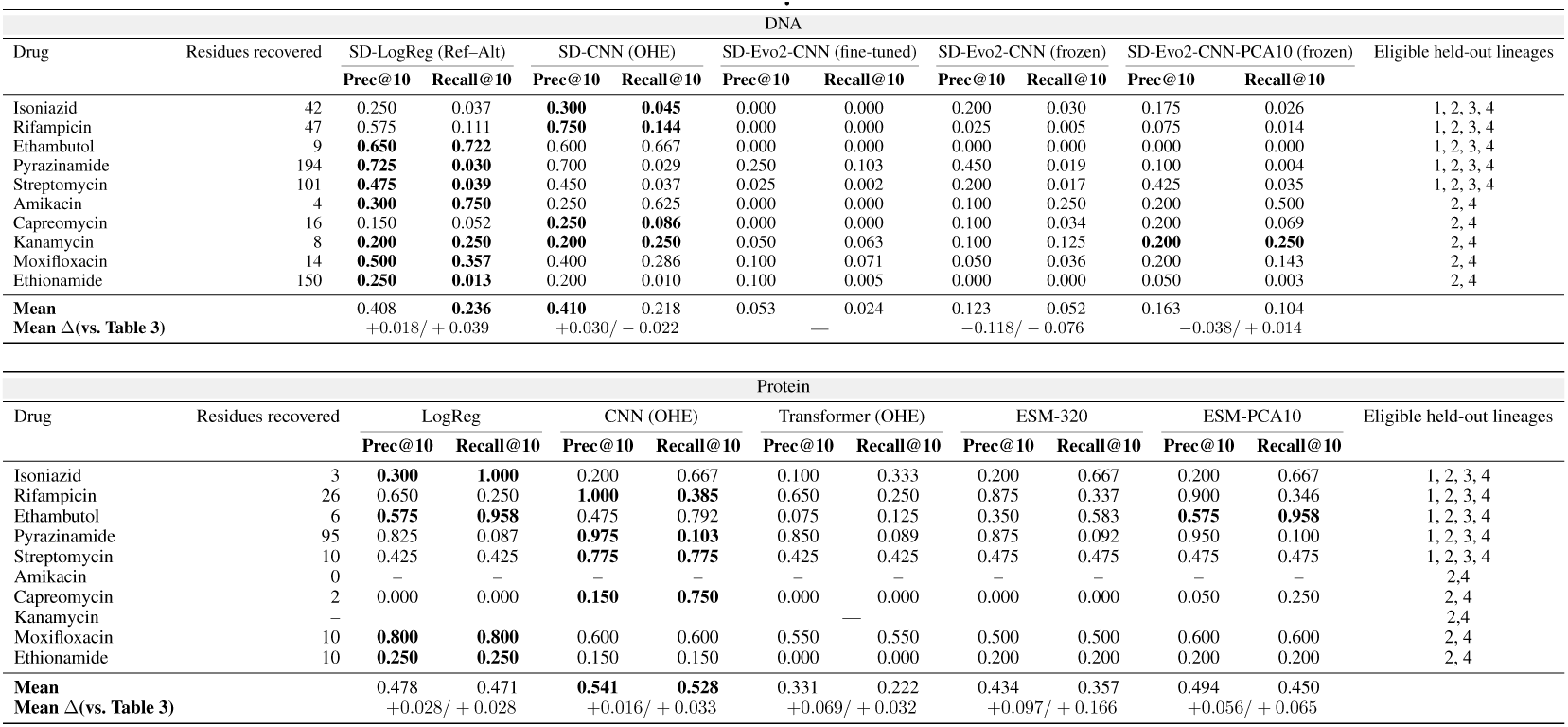
Precision@10 and Recall@10 for recovery of WHO-catalogued resistance target positions under lineage holdout, averaged over eligible held-out lineages (Section 4.5); “—” marks unavailable results and bold the best value per metric and drug within each modality. The **Mean** row averages each model over the drugs shown; **Mean** Δ is the difference from the random-spli.t values in Table 3

| DNA |  |  |  |  |  |  |  |  |  |  |  |  |
|---|---|---|---|---|---|---|---|---|---|---|---|---|
| Drug | Residues recovered | SD-LogReg (Ref–Alt) |  | SD-CNN (OHE) |  | SD-Evo2-CNN (fine-tuned) |  | SD-Evo2-CNN (frozen) |  | SD-Evo2-CNN-PCA10 (frozen) |  | Eligible held-out lineages |
|  |  | **Prec@10** | **Recall@10** | **Prec@10** | **Recall@10** | **Prec@10** | **Recall@10** | **Prec@10** | **Recall@10** | **Prec@10** | **Recall@10** |  |
| Isoniazid | 42 | 0.250 | 0.037 | **0.300** | **0.045** | 0.000 | 0.000 | 0.200 | 0.030 | 0.175 | 0.026 | 1, 2, 3, 4 |
| Rifampicin | 47 | 0.575 | 0.111 | **0.750** | **0.144** | 0.000 | 0.000 | 0.025 | 0.005 | 0.075 | 0.014 | 1, 2, 3, 4 |
| Ethambutol | 9 | **0.650** | **0.722** | 0.600 | 0.667 | 0.000 | 0.000 | 0.000 | 0.000 | 0.000 | 0.000 | 1, 2, 3, 4 |
| Pyrazinamide | 194 | **0.725** | **0.030** | 0.700 | 0.029 | 0.250 | 0.103 | 0.450 | 0.019 | 0.100 | 0.004 | 1, 2, 3, 4 |
| Streptomycin | 101 | **0.475** | **0.039** | 0.450 | 0.037 | 0.025 | 0.002 | 0.200 | 0.017 | 0.425 | 0.035 | 1, 2, 3, 4 |
| Amikacin | 4 | **0.300** | **0.750** | 0.250 | 0.625 | 0.000 | 0.000 | 0.100 | 0.250 | 0.200 | 0.500 | 2, 4 |
| Capreomycin | 16 | 0.150 | 0.052 | **0.250** | **0.086** | 0.000 | 0.000 | 0.100 | 0.034 | 0.200 | 0.069 | 2, 4 |
| Kanamycin | 8 | **0.200** | **0.250** | **0.200** | **0.250** | 0.050 | 0.063 | 0.100 | 0.125 | **0.200** | **0.250** | 2, 4 |
| Moxifloxacin | 14 | **0.500** | **0.357** | 0.400 | 0.286 | 0.100 | 0.071 | 0.050 | 0.036 | 0.200 | 0.143 | 2, 4 |
| Ethionamide | 150 | **0.250** | **0.013** | 0.200 | 0.010 | 0.100 | 0.005 | 0.000 | 0.000 | 0.050 | 0.003 | 2, 4 |
| **Mean** |  | 0.408 | **0.236** | **0.410** | 0.218 | 0.053 | 0.024 | 0.123 | 0.052 | 0.163 | 0.104 |  |
| **Mean Δ(vs. Table 3)** |  | +0.018/ + 0.039 |  | +0.030/ − 0.022 |  | — |  | −0.118/ − 0.076 |  | −0.038/ + 0.014 |  |  |
| Protein |  |  |  |  |  |  |  |  |  |  |  |  |
| Drug | Residues recovered | LogReg |  | CNN (OHE) |  | Transformer (OHE) |  | ESM-320 |  | ESM-PCA10 |  | Eligible held-out lineages |
|  |  | **Prec@10** | **Recall@10** | **Prec@10** | **Recall@10** | **Prec@10** | **Recall@10** | **Prec@10** | **Recall@10** | **Prec@10** | **Recall@10** |  |
| Isoniazid | 3 | **0.300** | **1.000** | 0.200 | 0.667 | 0.100 | 0.333 | 0.200 | 0.667 | 0.200 | 0.667 | 1, 2, 3, 4 |
| Rifampicin | 26 | 0.650 | 0.250 | **1.000** | **0.385** | 0.650 | 0.250 | 0.875 | 0.337 | 0.900 | 0.346 | 1, 2, 3, 4 |
| Ethambutol | 6 | **0.575** | **0.958** | 0.475 | 0.792 | 0.075 | 0.125 | 0.350 | 0.583 | **0.575** | **0.958** | 1, 2, 3, 4 |
| Pyrazinamide | 95 | 0.825 | 0.087 | **0.975** | **0.103** | 0.850 | 0.089 | 0.875 | 0.092 | 0.950 | 0.100 | 1, 2, 3, 4 |
| Streptomycin | 10 | 0.425 | 0.425 | **0.775** | **0.775** | 0.425 | 0.425 | 0.475 | 0.475 | 0.475 | 0.475 | 1, 2, 3, 4 |
| Amikacin | 0 | – | – | – | – | – | – | – | – | – | – | 2,4 |
| Capreomycin | 2 | 0.000 | 0.000 | **0.150** | **0.750** | 0.000 | 0.000 | 0.000 | 0.000 | 0.050 | 0.250 | 2, 4 |
| Kanamycin | – |  |  |  |  | — |  |  |  |  |  | 2,4 |
| Moxifloxacin | 10 | **0.800** | **0.800** | 0.600 | 0.600 | 0.550 | 0.550 | 0.500 | 0.500 | 0.600 | 0.600 | 2, 4 |
| Ethionamide | 10 | **0.250** | **0.250** | 0.150 | 0.150 | 0.000 | 0.000 | 0.200 | 0.200 | 0.200 | 0.200 | 2, 4 |
| **Mean** |  | 0.478 | 0.471 | **0.541** | **0.528** | 0.331 | 0.222 | 0.434 | 0.357 | 0.494 | 0.450 |  |
| **Mean Δ(vs. Table 3)** |  | +0.028/ + 0.028 |  | +0.016/ + 0.033 |  | +0.069/ + 0.032 |  | +0.097/ + 0.166 |  | +0.056/ + 0.065 |  |  |

Task 2 evaluates whether model-attributed positions recover WHO-catalogued resistance variants. It therefore measures agreement between model attribution and a curated reference set; it does not establish that a recovered site is the complete causal explanation for a resistance phenotype, nor that an unrecovered site is biologically irrelevant. A limitation of Task 2 is that the evaluation is primarily position-centric, whereas some resistance phenotypes depend on combinations of mutations across loci [38, 39] or compensatory changes [40]. BIG-TB can support evaluation with future interpretability methods, including mechanistic and causal approaches.

The lineage-held-out analysis provides important findings for the DNA and protein-based models. Mean ΔAUC for Task 1 phenotype prediction was small for all four DNA and all five protein architectures (DNA: *−*0.036 to *−*0.048; protein: *−*0.012 to *−*0.046,Supplementary Tables 18, 17), indicating that most of the predictive signal is preserved across held-out *M. tuberculosis* lineages and is not primarily driven by *M. tuberculosis* population structure, although these small declines were statistically significant for some models after correction.

BIG-TB provides a reusable benchmark for investigating the relationship between predictive performance, biological interpretability, and robustness in sequence-based antibiotic resistance prediction. Rather than evaluating model quality solely through phenotype prediction accuracy, BIG-TB enables researchers to examine whether improvements in predictive performance are accompanied by stronger recovery of established resistance-associated positions and whether these properties are maintained across distinct genetic backgrounds. Our evaluation illustrates the importance of these complementary objectives: models achieving higher resistance prediction accuracy did not necessarily recover more WHO-catalogued resistance positions, and pretrained sequence representations did not consistently outperform simpler supervised baselines. By providing curated data, reference variant annotations, baseline implementations, and reproducible evaluation procedures, BIG-TB offers a common experimental framework for developing and comparing future sequence-based models in pathogen genomics.

## 4 Methods

This section summarizes the data, tasks, models and evaluation design. Full details are given in the Supplementary Material: WHO catalogue processing and sequence curation in Supplementary Sections A–F, model architectures, training and SHAP implementation in Supplementary Section G, and additional methods (variant calling, embedding variants, the WHO heuristic and lineage-holdout splits) in Supplementary Section H.

### 4.1 Datasets

#### Phenotypes and genomes

We compiled laboratory-determined phenotypes for *M. tuberculosis* clinical isolates from previously published studies [41–45, 38, 46–52, 31, 53–56] for eleven antibiotics: rifampicin, isoniazid, ethambutol, pyrazinamide, streptomycin, capreomycin, amikacin, ethionamide, moxifloxacin, levofloxacin, and kanamycin. Raw reads were downloaded from the NCBI SRA and uniformly reprocessed: variants were called against the H37Rv reference [57] with an established pipeline [58, 59], and isolates with *<* 95% of the reference covered at 20*×* were removed (Supplementary Section H.1).

#### Sequence inputs

From each isolate’s VCF we reconstruct gene-centric DNA sequences for resistance-associated loci [60, 36] and align them into multiple sequence alignments (Figure 1; Appendices C and D). Protein sequences are obtained by translation with BioPython [61]; isolates with a frameshift in a locus are excluded from protein inputs for that locus (Appendix E). Our code supports arbitrary genomic regions, but the evaluations here are gene-centered and do not test whole-genome, zero-shot discovery of distal resistance mechanisms.

#### WHO resistance catalogue

We linked isolate genotypes to the WHO 2023 catalogue of resistance-associated mutations [30], which grades variants into five confidence groups (Supplementary Section H.3; Appendix B). Annotations are summarized in Table 1; Groups 1 and 2 (associated with resistance, including interim) form the Task 2 reference set.

### 4.2 Tasks

#### Task 1: phenotype prediction

Following prior work [62], models predict resistance (R) or susceptibility (S) from sequence, either for one drug (single-drug models, using that drug’s loci) or jointly for all eleven drugs (multi-drug models, using all loci; Table 8). Performance is measured by ROC-AUC.

#### Task 2: WHO-catalogued resistance varihant recovery

Following Tasmin and Green [35], we test whether model attributions recover WHO Group 1/2 variants. For each drug and modality, catalogue entries within the model input are collapsed to unique target positions. Sites are ranked by attribution magnitude, computed with DeepSHAP [16, 17] for neural models and the SHAP LinearExplainer for logistic regression (Appendix G.4), and compared with the target positions using Precision@*k* (fraction of the top-*k* sites that are target positions) and Recall@*k* (fraction of target positions among the top *k*), with *k* = 10 as the primary endpoint.

### 4.3 Models

We compare three encoding families: (i) reference–alternate encodings with logistic regression; (ii) one-hot encodings (DNA: 5 symbols including gaps; protein: 20 residues) with CNNs following Green et al. [36] and, for protein, Transformers; and (iii) foundation-model embeddings, Evo2-7B for DNA (*d* = 4096 per token) [7] and ESM-2 (esm2_t6_8M_UR50D, *d* = 320) for protein [63], extracted from the frozen pretrained model and used as input to shallow 1D CNNs of the same design. Because full per-token embeddings are memory-intensive, we also evaluate compressed variants (PCA-10, channel-mean, and sequence-mean; Supplementary Section H.4). For DNA, we additionally evaluate a multi-drug CNN (MD-CNN), LoRA fine-tuning of Evo2 under lineage holdout, and a non-learned WHO-catalogue heuristic that calls an isolate resistant if it carries any Group 1/2 variant for the drug (Supplementary Section H.6; Table 10). Architectures are given in Supplementary Section G.2 and Table 11, training hyperparameters in Supplementary Section G.3, input loci and how they are combined in Supplementary Section H.5, and SHAP settings in Supplementary Section G.4.

### 4.4 Training, validation, and test data

#### Unit of data and model-ready cohort

Each sample is one clinical isolate. For a given drug, the model-ready cohort comprises all isolates with a resistant or susceptible phenotype and an eligible (frameshift-free) sequence at every input locus for that drug (Section 4.3); for drugs with several loci, an isolate’s loci are concatenated into a single input. Phenotype class counts per drug are given in Table 1.

#### Held-out test set

For each drug, the model-ready cohort was split once into training (80%) and test (20%) partitions, stratified by phenotype (seed = 42). The split is performed at the isolate level, after concatenating loci, so each isolate appears in exactly one partition. Models were trained on the training partition only and evaluated once on the test partition; the test partition was not used for training, tuning, or model selection. Test AUCs from this split are reported in Table 2, and per-drug train/test counts in Table 14.

#### Cross-validation

Separately, we estimated per-drug performance with 5-fold stratified cross-validation (CV) over the full model-ready cohort (seed = 42), again with isolates as the unit, so that each isolate is in the validation fold exactly once and never in the training folds of the same split. CV models are distinct from the held-out-test models. We summarize CV ROC-AUC as mean *±* standard deviation across folds (Figure 2) and use the per-drug mean CV AUC for the model-vs-baseline significance tests (Section 4.6).

#### Sequence similarity between training and test isolates

Because *M. tuberculosis* is clonal and slowly evolving, isolates can share identical genotypes at the target loci. Identical genotypes in different isolates are distinct biological samples that can carry different phenotypes, so we do not remove them; instead, we assess generalization to genetically distinct backgrounds with the lineage-holdout evaluation(Section 4.5), in which all isolates from one major lineage are withheld from training and used as the test set.

### 4.5 Lineage-holdout evaluation

To assess sensitivity to *M. tuberculosis* population structure, we repeated Tasks 1 and 2 with lineage-holdout splits. Lineages were called with fast-lineage-caller and collapsed to major lineages 1–4. Each split withholds an entire major lineage (hundreds to thousands of isolates) as an independent test set and trains on all remaining isolates, so performance is measured on a genetic background absent from training; this tests robustness to population structure rather than performing leave-one-out cross-validation over individual isolates. A split was retained only if both partitions contained at least 50 resistant and 50 susceptible isolates, giving 30 drug/lineage settings for DNA (both tasks) and 28 (Task 1) and 26 (Task 2) for protein; levofloxacin had no eligible split (Supplementary Section H.7). Each lineage-holdout model was retrained from scratch with the random-split settings. For Task 2, the SHAP explain set was fixed to the isolates explained in the random-split analysis(Supplementary Section H.8). Among DNA foundation models, the lineage evaluation used Evo2, both frozen and LoRA fine-tuned.

### 4.6 Statistical testing across models

All significance tests are paired, two-sided Wilcoxon signed-rank tests in which the unit of pairing is the drug. Cross-validation folds are used only to estimate each drug’s performance: for each drug and model, the five fold AUCs are averaged to a single per-drug mean AUC, and per-fold values are shown in Figure 2.

We use this drug-level test for four comparisons: (i) *Task 1, model vs. baseline*: each model’s per-drug mean cross-validation AUC is paired with the Logistic Regression baseline’s per-drug mean cross-validation AUC (*n* = 11 drugs for DNA, *n* = 10 for protein; Tables 15 and 16); (ii) *Task 1, lineage holdout vs. random split*: each model’s per-drug lineage-holdout AUC, averaged over eligible held-out lineages, is paired with the same model’s random-split held-out test AUC (Table 2; *n* = 10 DNA, *n* = 9 protein); (iii) *Task 2, model vs. baseline*: each model’s per-drug Precision@10 and Recall@10 under the random split are paired with the Logistic Regression baseline’s values (*n* = 11 DNA, *n* = 10 protein); (iv) *Task 2, lineage holdout vs. random split*: each model’s per-drug Precision@10 and Recall@10 under lineage holdout are paired with the same model’s random-split values from the controlled-SHAP comparison (*n* = 10 DNA, *n* = 8 protein). Precision@10 and Recall@10 are tested separately.

Within each table and metric, we apply Holm–Bonferroni correction across the architectures included in that comparison, as specified in the corresponding table caption, following Demšar [64]. We report both raw and Holm-corrected *p*-values and use *p <* 0.05 after correction as the significance threshold. This framework was applied to the model families evaluated in each setting.

## Code and Data Availability

Code is available at https://github.com/SAGE-Lab-UMass/Big-TB-benchmark. Data are available at https://doi.org/10.6084/m9.figshare.c.8783478 The released benchmark provides reusable data-processing and evaluation components, enabling researchers to integrate additional sequence-based models while retaining the standardized phenotype prediction and WHO-catalogued resistance position recovery tasks. Evaluation procedures support comparisons under random and lineage-holdout splits, allowing newly developed models to be assessed against the baseline architectures reported here. The repository includes instructions for reproducing the benchmark results and applying the evaluation framework to additional models.

## Acknowledgments and Disclosure of Funding

The authors thank members of the UMass SAGE lab and the Farhat lab at HMS DBI for valuable input. This work utilized resources from Unity, a collaborative, multi-institutional high-performance computing cluster managed by UMass Amherst Research Computing and Data. SK was supported by the National Science Foundation Graduate Research Fellowship DGE2140743.

## Author Contributions

M.T. and S.M. contributed equally to this work. M.T., S.M., and A.G.G. conceived and designed the study. S.M. curated the genotype data and constructed the DNA sequence alignments. S.K. provided the variant call files and lineage-calling code. M.T. developed the protein-level models and performed the protein analyses. S.M. developed the DNA-level models and performed the DNA analyses. A.G.G. and M.R.F. supervised the study. M.T. led manuscript preparation and figure generation; all authors reviewed and edited the manuscript.

