## Supplementary Information for "BIG-TB: A benchmark for prediction and interpretability of sequence-based machine learning using *Mycobacterium tuberculosis* genomes"

### SUPPLEMENTARY MATERIALS

#### A Catalogue Overview

We downloaded the 2nd edition (2023) of the WHO Mutation Catalogue for *M. tuberculosis* complex and its association with drug resistance in Excel format (version *WHO-UCN-TB-2023.6-eng.xlsx*) [30]. The catalogue includes over 30,000 observed variants from 52,000 isolates worldwide, detailing the frequency of each mutation and its resistance or susceptibility associations, for 15 anti-TB drugs. We created a cleaned, user-friendly CSV version of this catalogue. Each row in the cleaned dataset represents a mutation and its corresponding resistance associations, with one row per mutation per drug. Our cleaned WHO catalogue has the following columns:

1. Variant: Describes the mutation, including the affected gene, the type of mutation (e.g., nucleotide change, amino acid change, or loss of function (`LoF`)), and its position, including any upstream or downstream regions. The notation follows the Human Genome Variation Society (HGVS) [65] guidelines. Examples of the variant column representations are shown below:
   - `bacA_c.1527C>T`: A nucleotide substitution in the bacA gene of the H37Rv reference genome, where cytosine (C) is replaced by thymine (T) at position 1527 in the gene's coding region.
   - `bacA_c.-21G>A`: A nucleotide substitution in the bacA gene of the H37Rv reference genome, where guanine (G) is replaced by adenine (A) 21 bases upstream of the start codon (ATG) in the untranslated region of the gene.
   - `bacA_p.Ala197Val`: An amino acid substitution in the bacA gene of the H37Rv reference genome, where alanine (Ala) is replaced by valine (Val) at position 197 in the protein sequence.
   - `bacA_LoF`: A Loss of Function (LoF) mutation in the bacA gene.
   - `rrs_n.1000G>A`: A nucleotide substitution in the rrs gene of the H37Rv reference genome, where guanine (G) is replaced by adenine (A) at position 1000 in the gene's non-coding RNA region of the sequence.

   The positions in this notation represent the relative positions with respect to the coding regions or start positions of the gene, unlike the genome index column, which represents an absolute position with respect to the H37Rv reference genome.
2. Drug: The antibiotic being considered. The catalogue includes 15 first-line drugs: isoniazid, rifampicin, pyrazinamide, and ethambutol, as well as second-line drugs: streptomycin, ciprofloxacin, delamanid, moxifloxacin, clofazimine, capreomycin, amikacin, kanamycin, ethionamide, bedaquiline, and linezolid.
3. Confidence: The confidence level assigned to the mutation in causing resistance to the drug under consideration. The confidence categories are:
   - 1) Assoc w R
   - 2) Assoc w R - Interim
   - 3) Uncertain significance
   - 4) Not assoc w R - Interim
   - 5) Not assoc w R

   Categories 1 and 2 are associated with resistance, while categories 4 and 5 are not associated and therefore consistent with susceptibility when observed. Category 3 indicates uncertainty regarding the association with resistance or susceptibility.
4. Genome_index: The starting index of the nucleotide subsequence on the forward strand of the H37Rv reference genome containing the mutation. For example, if `ATG` changes to `TGC` at position 32,467, this denotes the absolute position of `A` in the reference genome, marking the start of the mutated subsequence.
5. Gene: The gene in which the mutation has occurred.
6. Ref_nt: Refers to the original nucleotide subsequence in the H37Rv genome, before any mutation at the given genomic index.

7. Alt_nt: Refers to the alternate nucleotide subsequence observed as a mutation from the H37Rv genome, starting at the given genomic index.

The `ref_nt` and `alt_nt` represent nucleotide subsequences even for amino acid changes, as these correspond to mutations at the nucleotide level, regardless of their effect on the reference genome sequence.

The original WHO mutation catalogue consists of two sheets. The *Catalogue_master_file* contains columns for `Gene`, `Variant`, `Drug`, `Confidence`, and additional information. The *Genomic_coordinates* sheet includes columns for `Variant`, `Genome_index`, `Ref_nt`, and `Alt_nt` for all observed *M. tuberculosis* variants. First, we removed rows with missing values in the variant and position columns from the *Catalogue_master_file*. Entries with "combo" in the confidence column were excluded to ensure only clearly defined mutations remained. Then, we merged the sheets into a single dataframe using the variant column as the key. The dataframe includes all the aforementioned columns and was exported as *WHO_resistance_variant_all_2023.csv*.

Table 6: Distribution of resistance-associated variants in the WHO catalogue across all the confidence categories for 15 anti-TB drugs.

| Drug | Cat1 | Cat2 | Cat3 | Cat4 | Cat5 |
|---|---|---|---|---|---|
| Isoniazid | 3141 | 378 | 17089 | 3400 | 191 |
| Rifampicin | 95 | 358 | 15836 | 7933 | 337 |
| Ethambutol | 52 | 0 | 16926 | 4195 | 273 |
| Pyrazinamide | 1477 | 864 | 5035 | 1611 | 77 |
| Streptomycin | 795 | 586 | 8530 | 963 | 74 |
| Amikacin | 2 | 5 | 8294 | 620 | 396 |
| Capreomycin | 1208 | 338 | 10398 | 434 | 388 |
| Kanamycin | 13 | 2 | 8671 | 652 | 53 |
| Moxifloxacin | 37 | 25 | 6291 | 2646 | 129 |
| Levofloxacin | 43 | 19 | 7172 | 2860 | 146 |
| Ethionamide | 2045 | 1136 | 6486 | 950 | 10 |
| Linezolid | 7 | 7 | 3912 | 111 | 13 |
| Bedaquiline | 638 | 2044 | 7754 | 789 | 2 |
| Clofazimine | 606 | 1986 | 19148 | 1140 | 40 |
| Delamanid | 0 | 11143 | 2029 | 748 | 0 |

#### B Mutation–Phenotype Interpretability Dataset Construction

**Codon level mutation grouping in the VCF file** A VCF file records each nucleotide change individually, while the WHO mutation catalogue includes both nucleotide and amino acid level mutations. To correctly merge the VCF data with the WHO catalogue, nucleotide mutations affecting the same amino acid residue must be grouped together. This ensures nucleotide-level mutations in the VCF can be mapped to amino acid-level mutations in the WHO catalogue. The steps for combining codons in the VCF files are outlined below:

1. Valid SNPs are selected for codon-level combination by removing mutations with the `IMPRECISE` flag, excluding insertions (`IC`) and deletions (`DC`), and retaining only those with equal-length `REF` and `ALT` alleles (`len(ALT) = len(REF)`). Indels are retained for separate downstream processing.
2. Each mutation is mapped to its codon by first identifying its gene using a genome position-to-gene dictionary. If mutation is in a coding region, the gene's coordinates are used to locate the codon based on the reading frame. For overlapping genes, codons from all relevant genes are retrieved.
3. SNPs within the same codon are combined into a single entry by merging the `REF` and `ALT` nucleotides into one sequence. For non-consecutive mutations, missing positions are filled with the reference nucleotide, and the position is set to the first nucleotide of the codon. The resulting entries are exported to an updated VCF file named `{filename}_combined_codons.vcf`.

Table 7: Snapshot of mutations for one isolate in CSV format, showing annotations and isoniazid `INH` specific phenotype and confidence. These fields are drug-specific and are populated only when the mutation is listed for the respective drug in the WHO catalogue. The complete file for an isolate includes phenotypes for all 11 drugs.

| Mutation | Gene | Position | Mutation Type | INH | INH_Confidence |
|---|---|---|---|---|---|
| inhA_c.-154G>A | inhA | 1674048 | intergenic_variant | R | 1) Assoc. w R |
| Rv2752c_p.Glu142* | Rv2752c | 3065768 | stop_gained | N/A | 3) Uncertain significance |
| dnaA_c.1131C>A | dnaA | 4139246 | synonymous_variant | S | 4) Not assoc. w R - Interim |
| glpK_p.Val460Ala | glpK | 4138377 | missense_variant | S | 5) Not assoc. w R |
| rpsL_c.-165T>C | rpsL | 781395 | intergenic_region | | |

**VCF annotation.** The updated VCF files are annotated with additional mutation details for seamless integration with the WHO catalogue. We use the `snpEff` tool [66] to annotate each mutation with the gene in which it occurs, the resulting amino acid change due to nucleotide substitution(s), and the predicted mutation effect (e.g., missense, nonsense, frameshift). The annotated VCFs are then exported to a new file named `{filename}_combined_codons.eff.vcf`. Mutations affecting multiple genes are annotated with each gene's information and stored as separate records.

**Variant standardization and mapping.** Each annotated mutation in the VCF is assigned a standardized variant name based on the HGVS guidelines to match the WHO naming. This is achieved by combining the `gene` and `HGVS.c` or `HGVS.p` columns in the annotated VCF as `{gene}_{HGVS.c or p}`, depending on whether the mutation is at the protein level.

The annotated VCF and WHO catalogue files are mapped using the variant name, mutation position, reference allele, and altered allele, with separate mappings for missense and other variants. The final mapped file includes the following columns:

- Variant: The variant name in the HGVS format (e.g., `rpoB_p.Ser450Leu`).
- Gene: The gene where the mutation occurs.
- Mutation_position: The genomic index of the starting position of mutation in the H37Rv reference genome.
- Mutation_type: The effect of the mutation (e.g., `missense_variant`, `synonymous_variant`).
- Drug-specific confidence: Columns for each of the 11 anti-TB drugs, indicating the confidence category for the mutation against the drug.
- Drug-specific phenotype: Columns for each of the 11 anti-TB drugs, indicating whether the mutation confers resistance to the drug:
  - `R` for mutations with confidence `1) Assoc with R` or `2) Assoc with R - Interim`.
  - `S` for mutations with confidence `4) Not assoc with R - Interim` or `5) Not assoc with R`.
  - `N/A` for mutations with confidence `3) Uncertain significance`.

We also provide an API to apply stringent quality thresholds to each mutation in the annotated VCF file before mapping, to filter out low-confidence mutations. The threshold values are detailed in appendix section D.

### C Resistance-Conferring Gene Extraction

To manage the resource-intensive nature of our ML models, we select a subset of genes from the H37Rv genome. Following standard practice, we choose genes with putative resistance-conferring mutations. For each drug in the WHO Catalogue [30], we select genes with at least one mutation classified under confidence category `1) Assoc w R`. Operon coordinates were obtained from MycoBrowser [67] and extended $\pm 100$ bp to include potential regulatory regions. For genes that fall contiguously on the DNA strand, padding is applied only upstream or downstream to not repeat any regions from the subsequent gene. Example: As *gyrB* and *gyrA* are placed contiguously on the DNA strand, 100bp are added to only upstream of the *gyrB* strand and 100bp are added to only the downstream of the *gyrA* strand. Table 8 lists the genomic coordinates for each gene, the strand from which the original nucleotide sequence was extracted and direction in which the operon padding is applied.

Table 8: Padded start and end coordinates for each gene or operon region. Strand column depicts the strand from which the original gene specific nucleotide sequence was taken for further processing.

| Gene or Operon | Padded Start (genome) | Padded End (genome) | Strand | Padding Applied |
|---|---|---|---|---|
| `rrs` | 1471746 | 1473382 | pos | +100 upstream |
| `rrl` | 1473658 | 1476895 | pos | +100 downstream |
| `tlyA` | 1917840 | 1918846 | pos | +100 upstream and downstream |
| `embC` | 4239763 | 4243147 | pos | +100 upstream |
| `embA` | 4243233 | 4246517 | pos | none |
| `embB` | 4246514 | 4249910 | pos | +100 downstream |
| `ethA` | 4325904 | 4327473 | neg | +100 upstream |
| `ethR` | 4327549 | 4328299 | pos | +100 downstream |
| `fabG1` | 1673340 | 1674183 | pos | +100 upstream |
| `inhA` | 1674202 | 1675111 | pos | +100 downstream |
| `katG` | 2153789 | 2156211 | neg | +100 upstream and downstream |
| `eis` | 2714024 | 2715432 | neg | +100 upstream and downstream |
| `gyrB` | 5140 | 7267 | pos | +100 upstream |
| `gyrA` | 7302 | 9918 | pos | +100 downstream |
| `pncA` | 2288581 | 2289341 | neg | +100 upstream and downstream |
| `rpoB` | 759707 | 763325 | pos | +100 upstream |
| `rpoC` | 763370 | 767420 | pos | +100 downstream |
| `gid` | 4407428 | 4408302 | neg | +100 upstream and downstream |
| `rpsL` | 781460 | 782034 | pos | +100 upstream and downstream |

#### D Generating Multiple Sequence Alignments from VCF Files

The following sections describe the construction of high-quality multiple sequence alignments for each gene region using VCF files. Two alignment formats are generated: *aligned*, which includes gap characters to ensure all sequences for a gene region have the same length, and *unaligned*, which does not preserve the original sequence lengths and may vary across isolates for the same gene region.

**Reference Genome Region Extraction** We extract relevant genomic regions from the H37Rv reference genome (GenBank accession: `NC_000962.3` [57]) using the start and end positions of the selected gene regions. The positions within the gene regions are 0-indexed.

**Mutation Categorization and Quality Control Filtering** Each VCF file is sequentially processed to extract mutations, which are categorized into three types: `ref` for positions with no mutation (sequence matches the reference), `alt` for mutations where the reference nucleotide is replaced by an alternate, and `missing` for low-quality or ambiguous calls (e.g., marked as "N").
To retain only high-confidence mutations, we apply stringent quality filters based on the recommendations in [68]. Mutations are reclassified as `missing` if they fail any of the following thresholds: `QUAL` $> 10$ for variant call quality, `DP` $> 5$ to ensure sufficient coverage, `MQ` $> 30$ to avoid misaligned reads, and `BQ` $> 20$ for SNPs and MNPs to ensure reliable base support. Additionally, Mutations flagged as `Amb` in the `FILTER` field are classified based on allele frequency (`AF`): `AF` $> 0.75$ as `alt`, `AF` $< 0.25$ as `ref`, and `0.25 < AF < 0.75` as `missing`. Mutations with the `IMPRECISE` flag in the `INFO` field are also classified as `missing`.

**Mutation Introduction into the Reference Sequence** To generate an isolate genome, mutations from the VCF file are systematically applied to the corresponding H37Rv reference gene region. Mutation types, including SNPs and indels, are carefully introduced to ensure accurate alignment. The process involves the following steps:

1. Substitution-Based Mutation Handling: For Single Nucleotide Polymorphisms (SNPs), the reference (`REF`) and alternative (`ALT`) alleles are extracted from the VCF. If the mutation is categorized as `alt`, the reference base is replaced with the alternate base; if categorized as `missing`, it is replaced with `"N"`. The substitution is applied in place, so the sequence

length remains unchanged. For multi-nucleotide polymorphisms (MNPs), where multiple adjacent bases change, the entire affected region is replaced with the alternate sequence, while preserving sequence length.

2. Deletion Handling: Deletions remove nucleotides from the sequence, shortening it (`ALT` < `REF`). The affected reference nucleotides are identified and replaced with "`-`" to preserve alignment only in the *aligned* sequences. Large deletions extending beyond the defined genetic region are replaced entirely with gaps. If a deletion removes an entire gene or region, the affected sites are logged for reference.

3. Insertion Handling: Insertions introduce nucleotides not present in the reference, resulting in a longer sequence (`ALT` > `REF`). The inserted bases from `ALT` are added immediately after the reference position. If an insertion exceeds `100 bp`, it is replaced with "`N`" to avoid alignment issues. To preserve alignment, corresponding gap characters ("`-`") are inserted into the reference sequence only in the *aligned* sequences. A dictionary is maintained to record the positions within the gene where insertions occur and the number of nucleotides added. Before generating the final aligned sequences, these insertion sites are revisited to ensure all sequences remain the same length by padding shorter sequences with gap characters.

**FASTA Export** For each locus, the aligned sequences are exported to a FASTA file. The reference sequence is saved under the label `>MT_H37Rv`, and each isolate's mutated sequence is saved under its unique identifier. Additionally, a CSV file is generated for each gene region to record insertion site positions for easy tracking.

### E Protein Translation

The protein sequence generation pipeline starts by selecting alignment slices from the isolate FASTA files using the reference H37Rv coordinates. Then, nucleotide segments are extracted with non-gapped indices and translated into protein sequences using a gap- and ambiguity-tolerant function. We implement the following rules: codons with ambiguous bases (e.g., 'N') were translated to 'X', gaps ('-') leading to in-frame mutations were included, and sequences with frameshifts were flagged and handled appropriately for the downstream model.

We introduce an additional quality filter to ensure no unexpected behavior in protein translation: we require that the translated sequence length matches the expected reference protein length within a 5% margin of error, and that no internal stop codons (except at the terminal position) are present. Sequences failing these criteria are flagged as frameshifted and handled separately. For example, we found that for the *rpoB* dataset, `100%` of translations matched expected lengths.

### F Curated List of Protein-Substitution Variants from WHO Catalogue

We filter the WHO Catalogue Master File [30] to capture only protein-level substitutions, characterized by the format gene_p.Aaa123Bbb, where a reference amino acid is substituted at a specific position. For each variant, we parse the 3-letter amino acid codes, convert them to the corresponding 1-letter representation, and extract the mutation position to generate a the mutation label (e.g., p.A109C). This processing yields a clean dataset containing both the original variant notation and its 1-letter mutation equivalent, alongside associated drug, phenotype, and confidence information. The final catalogue is exported as a machine-readable CSV to facilitate downstream protein-level analysis of resistance-conferring mutations in *M. tuberculosis*.

### G Computational Pipeline and Models

#### G.1 Heuristic Model

Rules-based heuristic approaches are in regular use to predict resistance of *M. tuberculosis* from genome sequences [51, 69]. The heuristic model generates resistance predictions for each isolate using mutation-to-phenotype mappings (Table 7). We label an isolate as resistant if it carries at least

Table 9: Representative snapshot of WHO-listed resistance-associated protein variants across key M. *tuberculosis* genes.

| Gene | Drug | Variant | Effect | Confidence | Mutation |
|---|---|---|---|---|---|
| eis | Amikacin | eis_p.Trp36Arg | missense_variant | 3) Uncertain significance | p.W36R |
| embB | Ethambutol | embB_p.Met306Val | missense_variant | 1) Assoc w R | p.M306V |
| ethA | Ethionamide | ethA_p.Arg207Gly | missense_variant | 1) Assoc w R | p.R207G |
| gid | Streptomycin | gid_p.Ala134Glu | missense_variant | 1) Assoc w R | p.A134E |
| gyrA | Levofloxacin | gyrA_p.Asp94Gly | missense_variant | 1) Assoc w R | p.D94G |
| inhA | Isoniazid | inhA_p.Ser94Ala | missense_variant | 2) Assoc w R - Interim | p.S94A |
| katG | Isoniazid | katG_p.Ser315Thr | missense_variant | 1) Assoc w R | p.S315T |
| pncA | Pyrazinamide | pncA_p.His57Asp | missense_variant | 1) Assoc w R | p.H57D |
| rpoB | Rifampicin | rpoB_p.Ser450Leu | missense_variant | 1) Assoc w R | p.S450L |

Table 10: Heuristic model performance per drug. The columns represent total number of isolates (N), true positives (TP), false positives (FP), true negatives (TN), false negatives (FN), sensitivity, and specificity.

| Drug | N | TP | FP | TN | FN | Sensitivity | Specificity |
|---|---|---|---|---|---|---|---|
| Isoniazid | 17435 | 5275 | 222 | 6419 | 300 | 0.946 | 0.967 |
| Rifampicin | 17581 | 4612 | 281 | 11372 | 225 | 0.953 | 0.976 |
| Ethambutol | 15110 | 2333 | 802 | 11233 | 610 | 0.793 | 0.933 |
| Pyrazinamide | 12903 | 1711 | 339 | 7584 | 339 | 0.835 | 0.957 |
| Streptomycin | 7682 | 1768 | 405 | 4829 | 648 | 0.732 | 0.923 |
| Amikacin | 3705 | 492 | 56 | 2908 | 214 | 0.697 | 0.981 |
| Capreomycin | 3854 | 467 | 119 | 3060 | 207 | 0.693 | 0.963 |
| Kanamycin | 3576 | 577 | 60 | 163 | 10 | 0.983 | 0.731 |
| Moxifloxacin | 2868 | 262 | 97 | 255 | 16 | 0.942 | 0.724 |
| Levofloxacin | 269 | 69 | 18 | 175 | 7 | 0.908 | 0.907 |
| Ethionamide | 2153 | 543 | 421 | 505 | 82 | 0.869 | 0.545 |

one category 1 or 2 resistance mutation found in the WHO catalogue for the chosen drug; otherwise, it is labeled sensitive.

##### G.2 Neural Network Architectures

Table 11: Neural network architectures used across encoding types.

| Model | Layer / Operation | Dimensions |
|---|---|---|
| CNN (one-hot DNA / protein) | Input tensor | $(C \times L)$, where $C$=5 (DNA) or 20 (protein) |
| | Stem conv | $\text{Conv1d}(C, 64, 1)$ |
| | Conv block 1 | $\text{Conv1d}(64, 64, 12)$, ReLU, $\text{MaxPool1d}(3)$ |
| | Conv block 2 | $\text{Conv1d}(64, 32, 3)$, ReLU, $\text{Conv1d}(32, 32, 3)$, ReLU, $\text{MaxPool1d}(3)$ |
| | Fully connected | $\text{Linear}(\text{flat}, 256)$, ReLU, $\text{Linear}(256, 256)$, ReLU, $\text{Linear}(256, 1)$ |
| CNN (foundation embeddings) | Input tensor | $(d \times L)$, $d$=320 (ESM-2) or 4096 (Evo2) |
| | Stem conv | $\text{Conv1d}(d, 64, 1)$ |
| | Conv block 1 | $\text{Conv1d}(64, 64, 12)$, ReLU, $\text{MaxPool1d}(3)$ |
| | Conv block 2 | $\text{Conv1d}(64, 32, 3)$, ReLU, $\text{Conv1d}(32, 32, 3)$, ReLU, $\text{MaxPool1d}(3)$ |
| | Fully connected | $\text{Linear}(\text{flat}, 256)$, ReLU, $\text{Linear}(256, 256)$, ReLU, $\text{Linear}(256, 1)$ |
| Transformer (one-hot protein) | Input tensor | $(20 \times L)$ |
| | Input projection | $\text{Linear}(20, 128)$ + sinusoidal positional encoding |
| | Encoder | 2 TransformerEncoder layers ($d_{\text{model}}$=128, 4 heads, feed-forward 256, dropout 0.1) |
| | Classifier | Mean pooling over positions, $\text{Dropout}(0.5)$, $\text{Linear}(128, 1)$ |
| Mean-MLP (sequence-mean) | Input vector | $(d)$, $d$=320 (ESM) or 4096 (Evo2) |
| | Layers | $\text{Linear}(d, 128)$, ReLU, $\text{Dropout}(0.2)$, $\text{Linear}(128, 1)$ |

##### G.3 Model training

For all single-run experiments outside cross-validation, data were split 80/20 into training and test sets. Model hyperparameters are summarized in Section 4.4.

**Model training on one-hot encodings and frozen embeddings.** Protein CNN, Transformer and ESM-based classifiers were trained with Adam (learning rate $5 \times 10^{-4}$, batch size 32, up to 20 epochs). DNA SD-CNN and MD-CNN models used Adam (learning rate $1.23 \times 10^{-4}$, batch size 128) with drug-specific epoch counts. For Frozen Evo2-CNN, per-token representations were extracted from block 20 of the frozen Evo2-7B model (5,000 nucleotides per gene, concatenated across loci for multi-gene drugs), and CNN classifiers were trained with Adam (learning rate $5 \times 10^{-5}$, weight decay $1 \times 10^{-5}$, up to 30 epochs). Neural models used binary cross-entropy. Logistic regression baselines used balanced class weights. Full settings are in the code repository.

**Finetuning SD-Evo2-CNN model.** Evo2-7B was fine-tuned using low-rank adaptation (LoRA), with the pretrained backbone frozen and only the LoRA adapters and downstream CNN classifier updated. LoRA adapters with rank 8, scaling parameter 16, and dropout 0.1 were inserted into the three linear layers of each Evo2 MLP block up to and including block 20; all pretrained Evo2 parameters remained frozen. Token representations from block 20 were supplied to the downstream CNN classifier. Training used AdamW with learning rates of $1 \times 10^{-4}$ for the adapters and $1 \times 10^{-3}$ for the classifier, weight decay of 0.01, binary cross-entropy loss, and an effective batch size of 16 obtained through gradient accumulation. Models were trained for up to 10 epochs, with the best checkpoint selected by validation ROC-AUC using a minimum of three epochs and a patience of three further epochs before early stopping.

**Resource requirements for the models.** All models were implemented in Python. The SD-LogReg, SD-CNN, and MD-CNN models were implemented using TensorFlow v2.14 [70], whereas all DNA foundation and protein models were implemented in PyTorch [71] and scikit-learn [72]. Training and inference were executed on the UMass Unity Supercomputing Cluster. Each training job ran on NVIDIA A100 GPUs with 8 CPU cores and 8 GB of system memory for single-drug models, whereas multi-drug DNA models were allocated 100 GB of system memory. Foundation model embedding extraction for Evo2 and ESM was performed on high-memory GPUs with up to 40 GB of GPU memory.

**Resource requirements for frozen Evo2 embedding generation.** Frozen Evo2 token embeddings were generated independently for each resistance-associated locus using Evo2-7B, with each locus represented using a maximum sequence length of 5,000 bases. Each locus-level embedding-generation job used one NVIDIA A100 GPU with 80 GB of GPU memory and 12–20 CPU cores sharing up to 500 GB of system RAM. The observed peak memory usage varied substantially across loci, ranging from approximately 112 GiB to the 500-GiB allocation limit. Because some jobs reached this limit, 500 GB was retained as the maximum allocation memory. Most locus-level embedding jobs required approximately 3.1–4.3 GPU-hours, corresponding to equivalent per-drug computational costs of approximately 3.4–11.0 GPU-hours depending on the number of associated loci.

**Resource requirements for frozen Evo2 downstream modeling.** Each frozen Evo2-CNN downstream cross-validation fold used one NVIDIA A100-SXM4 GPU with 80 GB of GPU memory, eight CPU cores sharing upto 300-GB of system RAM. A single fold therefore required one GPU, whereas executing all five random-split folds concurrently required five GPUs. Peak system-memory usage varied substantially across drugs, ranging from approximately 12 GiB to the 300-GiB allocation limit per fold. Several of the drugs with larger genes or gene combinations like ethambutol, rifampicin, isoniazid and few others reached this limit hence, a 300-GB system-memory allocation per fold was therefore retained across drugs. Observed mean runtime per fold ranged from approximately 1 to 4 hours, corresponding to accumulated five-fold computational costs of approximately 1–20 GPU-hours across drugs.

**Resource requirements for Evo2 fine-tuning.** Evo2-7B was fine-tuned with LoRA separately for each drug and held-out lineage. Each fine-tuning job used one NVIDIA A100-SXM4 GPU with 80 GB of GPU memory and eight CPU cores. Before launching the finetuning jobs, we ran memory profiling runs that measured a peak GPU-memory usage of approximately 49.6 GB, and an 80-GB A100 was therefore used to provide sufficient memory headroom. Additionally 64 GB CPU RAM per job was provided a conservative margin for the runs. One GPU was required for each held-out-lineage job, with up to four GPUs used concurrently when four lineage-specific jobs for a drug were executed in parallel. The observed GPU time per completed lineage run ranged from approximately 1.1 to 44.0 hours across drugs.

##### G.4 Explaining sequence models with SHAP

**Motivation and design principles.** SHAP's DeepExplainer requires a background distribution; too small yields unstable estimates, too large becomes computationally prohibitive. To ensure coverage without duplication, we deduplicate all isolates and partition into disjoint sets: a background (10%, capped at 160) and an explained set (90%). This guarantees that each unique sequence is explained exactly once and never used as its own reference. Splits are stratified by resistance label (at least one R and one S if both exist), seeded, and logged for reproducibility. Our attribution design balances (i) faithfulness to the trained model, (ii) biological fidelity by covering all unique isolates including rare variants, and (iii) computational practicality given high-dimensional embeddings.

**Deduplication and disjoint split.** Let $\mathcal{D} = \{(x_i, y_i)\}_{i=1}^{N}$ be the deduplicated pool with $y_i \in \{0, 1\}$. We sample without replacement to form background indices $B$ (size $m$) and set the explained indices $E = \{1, \ldots, N\} \setminus B$. We use stratification to ensure both R and S must appear in $B$ when available. Default parameters are $\alpha = 0.10$, $B_{\max} = 160$, and $m_E^{\min} = 1$. An implementation sketch is provided below.

```
def disjoint_bg_explain_indices(N, bg_frac=0.10, max_bg=160, seed=42):
    rng   = np.random.default_rng(seed)
    perm  = rng.permutation(N)
    bgn   = int(round(bg_frac * N))
    bgn   = min(max(bgn, 1), N-1, max_bg)
    bg    = perm[:bgn]
    ex    = perm[bgn:]
    return bg, ex
```

**Neural models (CNNs over embeddings).** We apply SHAP `DeepExplainer` to trained CNNs over ESM embeddings. Models are wrapped to output $(B, 1)$ tensors, and SHAP values are computed on the explained set. Attributions are aggregated across classes and sliced into gene segments using known per-gene padded lengths.

**Linear models.** For logistic, ridge, and lasso classifiers, we use SHAP's `LinearExplainer` with two maskers: *Independent* (interventional, "true to the model") and *Impute* (correlation-aware, "true to the data"). This allows comparison with CNN-based models under the same discovery metric. Scores are obtained for each residue in each gene by the maximum absolute attribution across explained isolates.

**Post-processing and evaluation.** For each gene, residues are ranked by their maximum absolute SHAP value across explained isolates. Recovery is benchmarked against the WHO catalogue, restricted to entries labeled "Assoc w R" or "Assoc w R – Interim" with `intersectional=True`. WHO positions are 1-indexed and were converted to 0-indexed residue positions by subtracting 1. We report Precision@k, Recall@k, and F1@k per drug.

```
# Build deduplicated pool of train U test
bg_idx, ex_idx = disjoint_bg_explain_indices(N, bg_frac=0.10, max_bg=150)
background = X[bg_idx]; explained = X[ex_idx]

explainer = shap.DeepExplainer(model, [background])
phi = explainer.shap_values([explained], check_additivity=False)[0]

# Aggregate per-residue and slice by gene
for gene, (s,e) in gene_slices.items():
    scores = np.abs(phi[:, s:e]).max(axis=0)
    evaluate_topk(scores, WHO_sites[gene])
```

**Defaults and sensitivity.** We fix the background fraction at 10%, cap at 160 sequences, and use a fixed seed. Increasing background size beyond 160 yields marginal differences in rankings but substantially increases runtime. All splits and metadata (#background, #explained, seed) are logged for reproducibility.

##### G.5 Evo2 Embedding Variants

Evo2 produces 4096-dimensional token embeddings. For sequences padded or truncated to $L = 5000$, a single embedding requires approximately 41 MB in `float16`, or approximately 492 MB for a batch of 12. To reduce memory overhead, embeddings were stored using chunked memory-mapped files. We evaluated full token-level, PCA-10, channel-mean, and sequence-mean representations.

##### G.6 ESM Embedding Variants

A single protein sequence of length $L = 800$ requires approximately 0.5 MB of memory when stored in `float16` precision, corresponding to roughly 64 MB for a batch size of 128 sequences. To mitigate this bottleneck, embeddings are chunked and stored using memory-mapped files with similar unified training interface as the DNA data.

### H Additional Methods

#### H.1 Variant calling and quality control

For isolates with published phenotype data, we compile publicly available whole-genome sequences for *M. tuberculosis*. We downloaded the raw read data from the NCBI SRA and performed our own variant calling to ensure consistency across datasets generated by multiple groups. We perform variant calling against the H37Rv reference genome [57] and variants were called according to a pipeline introduced in Ezewudo et al. [58] and updated in Freschi et al. [59]. We trim and filter reads with FASTP [73], remove contaminated isolates using Kraken2, and align to the reference using BWA-MEM2 [74]. We remove duplicate reads using Picard (http://broadinstitute.github.io/picard/), and we drop isolates with <95% coverage of the reference genome at 20× coverage. This results in one Variant Call Format (VCF) file per isolate that contains one row per mutation observed relative to the reference sequence.

#### H.2 Sequence extraction and protein translation

Both foundation models and standard machine learning models for antibiotic resistance prediction use DNA or protein sequences as input. To support this, we provide a custom pipeline that outputs FASTA-formatted sequences from VCF data for each isolate as shown in Figure 1 (Appendix D). For models that take unaligned sequences as input, the sequences are generated without gaps. For downstream modeling in this study, we follow standard practice [60, 36] and select only genomic regions suspected to be involved in antibiotic resistance (Appendices C and D). While we have hand-selected regions for downstream modeling, we note that our code supports extraction of arbitrary genomic regions. The current benchmark evaluations are therefore gene-centered and do not test whole-genome, zero-shot discovery of distal resistance mechanisms.

For models trained on protein data, DNA sequences for each gene region are translated into protein sequences using BioPython [61], as shown in Figure 1 (Appendix E). Translations with an internal stop codon or a greater than 5% length discrepancy from the reference were flagged and excluded from protein-model inputs; no released model-ready protein sequence met this criterion. Table 12 shows the per-isolate DNA sequence, protein sequence, phenotype label, and frameshift flag retained in these tables.

Table 12: Representative examples of *rpoB* mutations from isolates with distinct primary mutation sites. Each row lists the isolate ID, phenotype for a representative drug (RIF), the start of the full gene sequence, and the corresponding DNA and protein windows centered on the mutation site. Only the first detected mutation per isolate is reported for display.

| Filename | Phenotype | DNA Sequence | Protein Sequence | Mutation Pos. |
|---:|:---:|---|---|---|
| SAMN02585979 | R | ACAAGC... | HKRRLSALGP... | Q429H |
| SAMEA2533629 | S | ACAAGC... | HKRRLSALGL... | P454L |
| SAMEA3558815 | R | CCCAGC... | PQRRLSALGP... | L443- |
| SAMN08436285 | S | ACAAGC... | HKRRLSALGP... | E344Q |
| SAMN03648257 | S | ACAAGC... | HKRRLSALGP... | D270E |

#### H.3 WHO catalogue grading

The WHO 2023 catalogue compiles observed *M. tuberculosis* mutations and their association with antibiotic resistance using matched whole-genome sequencing and phenotypic drug-susceptibility data from multiple settings [30]. Variants are graded using statistical evidence into five groups: (1) associated with resistance, (2) associated with resistance—interim, (3) uncertain significance, (4) not associated with resistance—interim, and (5) not associated with resistance. We merge this catalogue with isolate-level VCF genotypes using a custom pipeline (Appendix B) and summarize the resulting annotations in Table 1. For Task 2, Group 1 and Group 2 entries form the WHO-catalogued resistance variant reference set; a curated subset of protein substitutions reformatted for downstream use is shown in Table 9.

##### H.4 Embedding representation strategies

For both DNA (Evo2) and protein (ESM) models, the starting point is a per-token embedding matrix $\mathbf{H} \in \mathbb{R}^{L \times d}$, where $L$ denotes sequence length and $d$ the embedding dimension. While these full embeddings retain maximal information, they are memory-intensive to store for tens of thousands of isolates. Therefore, we evaluate three representation strategies with embeddings (Table 13): (i) PCA-$k$: project per-token embeddings onto the top principal components ($k = 10$ for DNA, $k \in \{10\}$ for protein); (ii) channel-mean: average across embedding channels per token, yielding a 1D descriptor per residue; and (iii) sequence-mean: average across the length dimension to obtain a single $d$-vector per sequence. The latter is widely used in NLP-based baselines and past ESM-1v benchmarks [12] but discards positional structure entirely [75, 76]. For DNA models, we tested all four variants in single-drug CNNs, and refrained from testing multi-drug models due to GPU memory limits. For protein models, We tested all four variants across drugs. We note that sequence-mean cannot be used for WHO-catalogued resistance variant recovery because it discards residue-level structure.

Table 13: Embedding Representation Strategies grouped by granularity. Per-site methods yield token/residue detail; per-sequence methods yield global summaries.

| Variant | Output Shape | Compression | Description |
|---|---|---|---|
| Per-site embeddings (retain token-level detail) | | | |
| Full-$d$ | $L \times d$ | None | Raw per-token embeddings. ($d = 4096$ for Evo2 7b, $d = 320$ for ESM). |
| PCA-$k$ | $L \times k$ | PCA | Incremental PCA on all tokens, project to top $k$ components. |
| Channel-mean | $L \times 1$ | Mean | Average across $d$ channels per token. |
| Per-sequence embedding (global descriptor) | | | |
| Sequence-mean | $1 \times d$ | Mean | Average across length dimension; discards positional detail. |

##### H.5 Architectures and input loci

Logistic regression on reference–alternate encoded sequences to provide a simple and interpretable linear baseline. We train convolutional models on one-hot encoded sequences following Green et al. [36] for DNA-based models and implementing a similar architecture for protein-based models. For foundation models, we use shallow 1D CNNs with the same architecture, but with input channels corresponding to the embedding dimension ($d$=320 for ESM-2 or $d$=4096 for Evo2). For complete layer dimensions, see Table 11.

For single-drug models, input loci are fused by concatenation along the sequence axis. For the multi-drug DNA one-hot CNN model, input loci are stacked along the channel dimension.

##### H.6 WHO catalogue heuristic baseline

As a non-learned, rules-based comparator for DNA-level phenotype prediction, we additionally evaluate a WHO-catalogue-derived heuristic: an isolate is labeled resistant to a given drug if its VCF contains at least one WHO 2023 category 1 (associated with resistance) or category 2 (associated with resistance—interim) mutation for that drug, and susceptible otherwise. This rule mirrors the mutation-lookup logic underlying widely used TB resistance-prediction tools such as Mykrobe and TBProfiler [77, 78], and provides a clinically interpretable, non-learned reference point (Table 10).

##### H.7 Lineage-holdout splits

To assess sensitivity to *M. tuberculosis* population structure, we added a lineage-aware robustness evaluation, applied to both Task 1 (phenotype prediction) and Task 2 (WHO-catalogued resistance variant recovery). We used isolate-level lineage annotations generated with fast-lineage-caller and collapsed hierarchical lineage calls to the top-level lineage assignment, so that sublineages such as 4.2 or 4.2.1 were treated as lineage 4 . Major lineages 1–4 were considered as candidate held-out test groups. Five well-powered drugs (rifampicin, isoniazid, ethambutol, pyrazinamide, and streptomycin) supported evaluation across all four major held-out lineages, whereas five sparser drugs (amikacin, capreomycin, kanamycin, moxifloxacin, and ethionamide) satisfied the minimum class-count criterion only for held-out lineages 2 and 4. This yields 30 reportable drug/held-out-lineage split settings for DNA models in both Task 1 and Task 2. Protein models have 28 settings for Task 1, because kanamycin's resistance locus is not protein-coding, and 26 for Task 2, because amikacin additionally has no protein-coding WHO-catalogued target position (Section H.8).Levofloxacin did not satisfy the split criterion for any held-out major lineage and was therefore excluded from this analysis.

For the lineage-aware experiments, we restricted the foundation-model analysis to Evo2. We evaluated both the frozen representation and a LoRA-fine-tuned Evo2 model to determine whether task-specific adaptation improved generalization to held-out lineages.

For each drug task, we constructed lineage-holdout splits using major lineages 1, 2, 3, and 4 as candidate held-out test groups. Each split withholds an entire major lineage (hundreds to thousands of isolates) as an independent test set; this is a group-level out-of-distribution evaluation, not leave-one-out cross-validation over individual isolates. For a given held-out lineage, the test set consisted of all model-ready isolates assigned to that lineage, while the training set consisted of all remaining lineage-annotated isolates. Thus, isolates assigned to non-major lineage labels were retained in training whenever they were not part of the held-out test group. A held-out lineage split was included only when both the training and test partitions contained at least 50 resistant and 50 susceptible isolates

The same isolate-level train/test assignments were applied across all model families for a given drug and held-out lineage, and each lineage-held-out model (one per drug, model family, and held-out lineage) was trained from scratch using the same architecture, hyperparameters, and training procedure as the corresponding random-split model.

##### H.8 Task 2 under lineage holdout: fixed explain sets

We repeated Task 2 using the lineage-holdout models described above, so that the only change relative to the original random-split Task 2 evaluation (Section G.4) is which isolates a model was trained on.

Isolating a genuine training-distribution effect from an artifact of explaining a different set of isolates requires holding the explain set fixed. We therefore fixed the SHAP explain set, for each drug and architecture, to the exact isolate identities used in that architecture's original random-split (Table 3) SHAP computation, recovered deterministically from the corresponding cached dedup/split indices and random seed ($\text{seed} = 42$). SHAP background, in contrast, was drawn from each lineage-held-out model's own training partition – excluding any isolate that overlapped the fixed explain set, so that background and explain sets remained disjoint – since the background distribution must reflect what that specific model was actually trained on rather than the full random-split cohort. Background size followed the same $10\%$/capped-at-160 convention as the random-split design (Section G.4) applied to this smaller, lineage-restricted training pool, and could therefore shrink below this target for drugs and lineages with fewer available training isolates. With the explain set held constant across the random-split and lineage-held-out conditions, any difference between the two Precision@$k$/Recall@$k$ values for a given drug and architecture reflects a genuine effect of training on fewer lineages rather than an artifact of the explain set itself.

Task 2 lineage results are reported for the same feasibility-eligible drug/lineage combinations as Task 1 (Section H.7): the five well-powered drugs across all four held-out lineages, and capreomycin, moxifloxacin, and ethionamide across the two lineages (2 and 4) for which they are feasible. Amikacin is excluded from the protein-modality Task 2 lineage evaluation because it has no protein-coding WHO-catalogued target position in *eis* (Table 3); kanamycin is excluded from all protein-modality Task 2 evaluation because its resistance locus is not protein-coding.

#### I Results and Statistical Significance

##### I.1 Ablation Study of Foundation Model Embedding Variants

We systematically evaluate alternative representation strategies for compressing and aggregating token-level embeddings from Evo2 (DNA) and ESM–2 (protein). These representations differ in the degree to which they preserve positional detail versus global context (Table 13). Specifically, we compare: (*i*) full per-token embeddings that retain all token-level and contextual information; (*ii*) PCA-$k$ projections that reduce dimensionality while maintaining token-level information; (*iii*) channel-mean embeddings that collapse across channels per position; and (*iv*) sequence-mean embeddings that pool across the entire sequence, producing a single vector descriptor. Sequence-mean embeddings are widely used in NLP for efficiency [76], but they discard positional structure and are therefore unsuited for residue-level interpretability. Exact per-drug $\Delta$AUC values for each embedding variant are shown in Figures 5 and 6, which complement the main grouped bar plots by visualizing the full performance matrix.

**Predictive performance (AUC).** For DNA models, the evaluated Evo2 representation strategies consistently improve predictive performance over the seq-mean (global pooling) baseline (Figure 4a). Channel-mean achieves the highest AUC in 6/11 drugs and full-token in 5/11, whereas PCA-10 and seq-mean never lead. Together, these results show that the choice of embedding representation substantially affects downstream performance.

For protein models, preserving per-residue structure while compressing channels via PCA consistently improves accuracy over the seq-mean (global pooling) baseline (Figure 4b). Across the ten drugs, PCA-10 outperforms seq-mean in eight cases (mean $\Delta\text{AUC} = +0.048$), while full-320 improves in seven ($+0.025$ on average). Channel-mean aggregation is consistently harmful ($-0.24$). Within-ESM comparisons show that PCA-10 achieves the highest AUC in 5/10 drugs, full-320 in 4/10; seq-mean and channel-mean never lead. Exact normalized $\Delta$AUC values are provided in Figure 6.This trend indicates that moderate compression (PCA-10) effectively preserves the discriminative content of full embeddings while improving stability across drugs.

**WHO-catalogued resistance variant recovery.** For DNA models, full-token and PCA-10 Evo2 representations achieve comparable recovery of WHO-catalogued resistance positions (Table 3), while sequence-mean is excluded because it removes positional structure. For protein models, PCA-10 and full-320 ESM embedding representations achieve competitive recovery of resistance-conferring residues (Table 3). Sequence-mean is again excluded from the evaluation due to lack of positional structure. Overall, these ablations reveal that retaining per-token positional structure—either through full or PCA-compressed embeddings—is critical for capturing WHO-catalogued resistance variants: while global sequence pooling retains enough information for accurate classification, it fails to localize WHO-catalogued resistance variants because spatial context is lost.

##### I.2 Lineage-holdout robustness

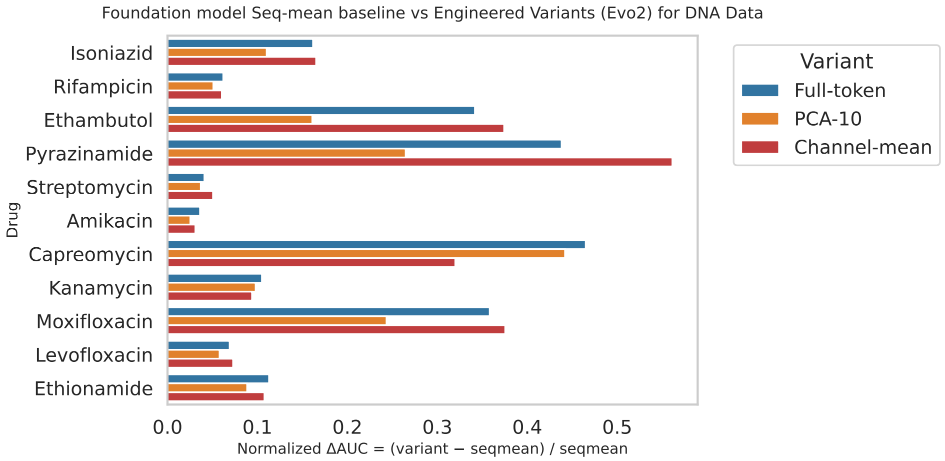


(a) Evo2 embedding representation strategies (DNA). Full-token representations outperform seq-mean across all 11 drugs (mean normalized $\Delta$AUC $= +0.199$), while PCA-10 also improves across all 11 drugs ($+0.144$ on average). Channel-mean aggregation similarly improves across all drugs and achieves the largest mean gain ($+0.201$). Within the Evo2 representations, channel-mean achieves the highest AUC in 6/11 drugs and full-token in 5/11, while PCA-10 and seq-mean never lead.

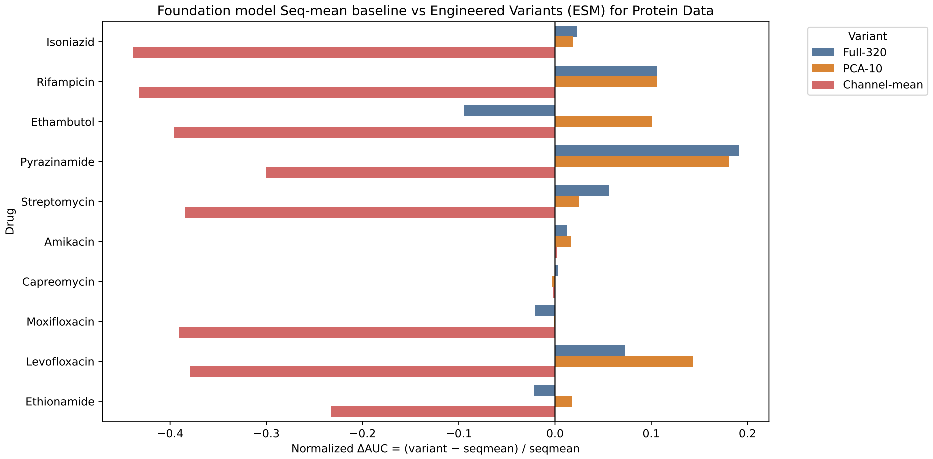


(b) ESM-2 embedding representation strategies (Protein). PCA-10 achieves consistent gains across 8/10 drugs (mean $\Delta$AUC $= +0.048$), with notable improvements for pyrazinamide ($+0.181$), levofloxacin ($+0.144$), rifampicin ($+0.106$), and ethambutol ($+0.100$). Full-320 provides smaller but stable gains ($+0.025$ on average), while channel-mean aggregation is consistently detrimental ($-0.24$).

Figure 4: Ablation of foundation model embedding representation strategies for DNA and protein sequences. Normalized change in AUC ($\Delta$AUC) is shown for each drug relative to the sequence–mean baseline. Positive values indicate improvement over seq-mean. Both Evo2 (a) and ESM (b) models show that preserving per-token structure (Full or PCA-10) yields clear accuracy gains, whereas channel-mean and global pooling substantially reduce predictive power. Exact normalized $\Delta$AUC matrices are provided in Figures 5,6

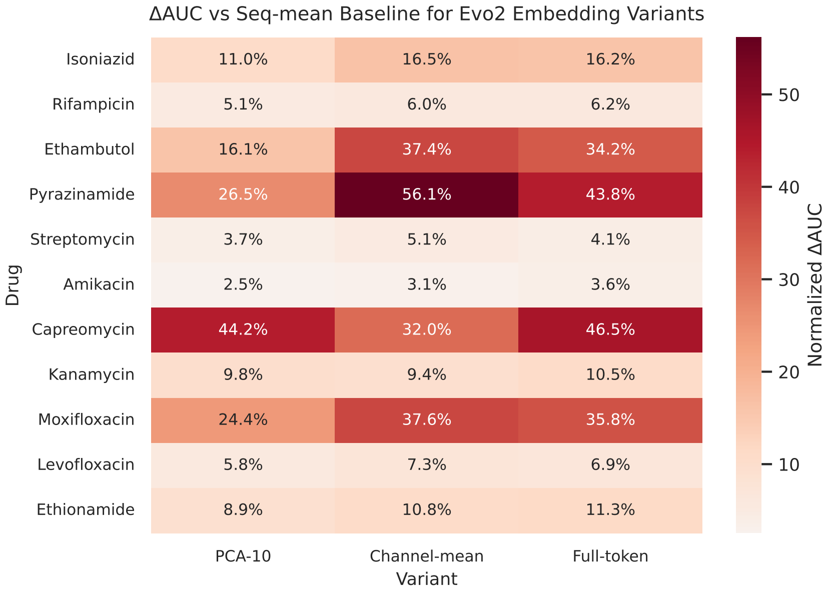


Figure 5: ΔAUC for Evo2 embedding representations. Full-768 and PCA-10 embeddings show consistent gains.

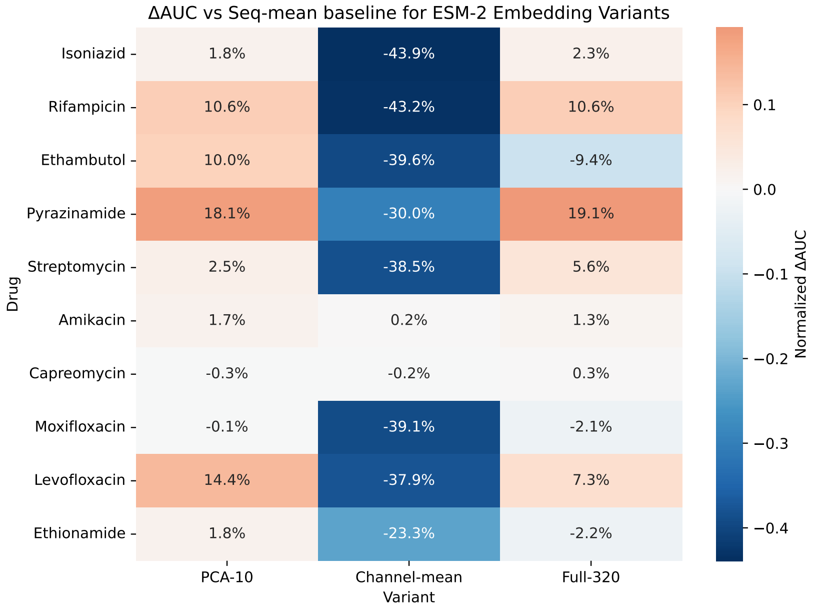


Figure 6: ΔAUC for ESM embedding representations. PCA-10 yields the largest gains for key drugs.

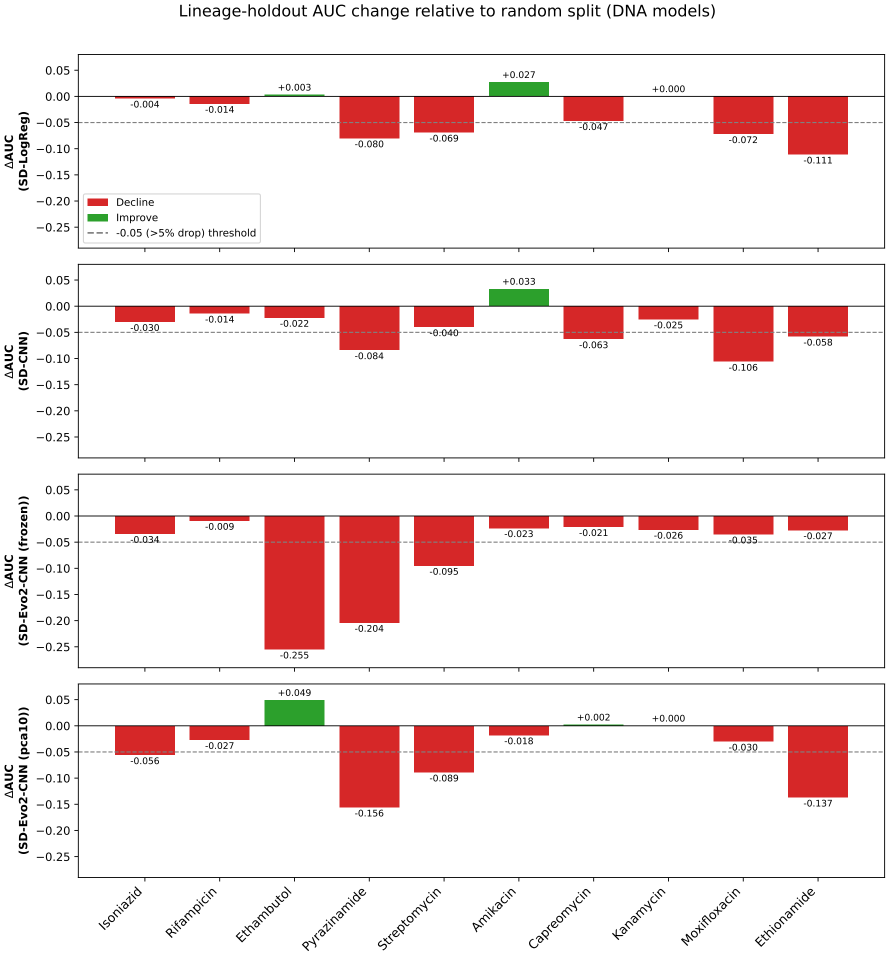


Figure 7: Per-model, per-drug lineage-holdout AUC change ($\Delta$AUC = lineage-split AUC minus random-split AUC, Table 2) for four DNA models: SD-LogReg, SD-CNN, SD-Evo2-CNN (frozen), and SD-Evo2-CNN-PCA10 (frozen). This complements the aggregated **Mean $\Delta$AUC** in Table 4 by highlighting architecture-specific effects. For example, ethambutol shows a sharp decline for frozen Evo2-CNN ($-0.255$) but an improvement for Evo2-CNN-PCA10 ($+0.049$), while LogReg ($+0.003$) and SD-CNN ($-0.022$) remain relatively stable. Pyrazinamide exhibits substantial degradation across all models, particularly frozen Evo2-CNN ($-0.204$) and Evo2-CNN-PCA10 ($-0.156$), compared with LogReg ($-0.080$) and SD-CNN ($-0.084$). Streptomycin similarly shows larger declines for both Evo2 variants ($-0.095$ and $-0.089$) than for LogReg ($-0.069$) and SD-CNN ($-0.040$), whereas amikacin improves for LogReg ($+0.027$) and SD-CNN ($+0.033$) but declines slightly for both Evo2 variants. The dashed line marks an AUC decline of 0.05; negative and positive values indicate degraded and improved performance, respectively.

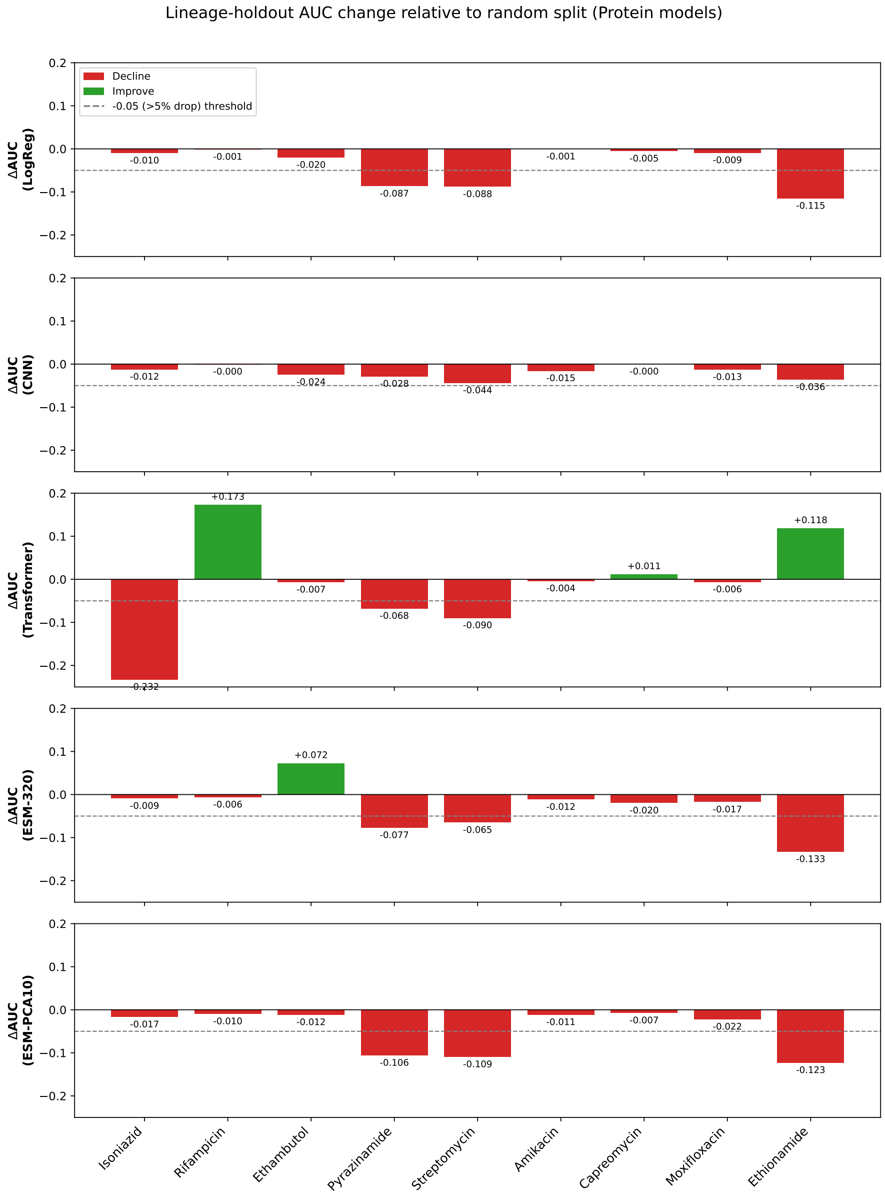


Figure 8: Per-model, per-drug lineage-holdout AUC change for the five protein models (LogReg, CNN, Transformer, ESM-320, ESM-PCA10), analogous to Figure 7. This complements the blended per-architecture **Mean ΔAUC** row in Table 4 to highlight large, partially offsetting architecture-specific effects: rifampicin's blended $+0.031$ mean across the five architectures is driven almost entirely by Transformer's $+0.173$ swing (the other four architectures are each within $\pm 0.010$), isoniazid's blended $-0.056$ mean is likewise driven by Transformer's $-0.233$ decline despite the other four architectures changing by at most 0.017, and ethambutol's near-zero blended mean ($+0.002$) masks ESM-320 improving by $+0.072$ while LogReg, CNN, Transformer, and ESM-PCA10 all decline modestly.

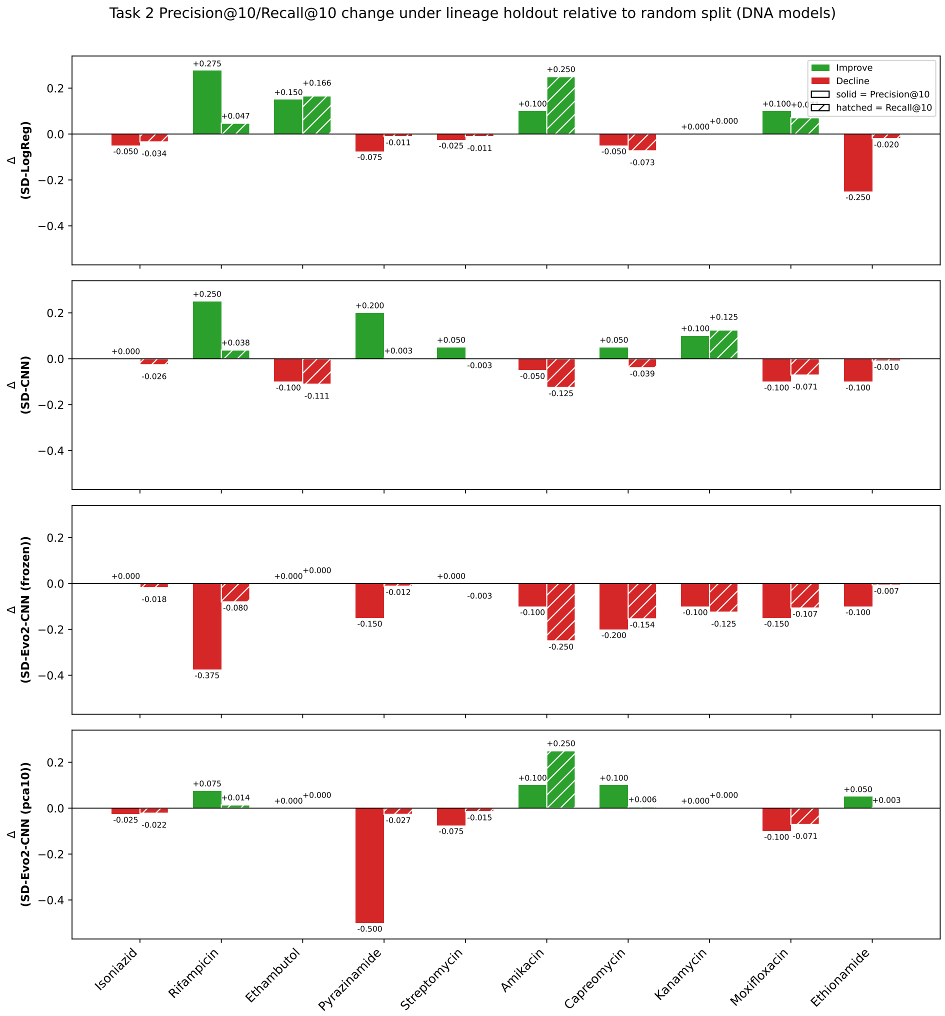


Figure 9: Per-model, per-drug change in Task 2 Precision@10 (left, solid bar) and Recall@10 (right, hatched bar) under lineage holdout relative to the random split (Table 3), for four DNA models: SD-LogReg, SD-CNN, SD-Evo2-CNN (frozen), and SD-Evo2-CNN-PCA10 (frozen). This complements the lineage-holdout results in Table 5, highlighting model-specific differences in recovering known resistance-associated mutations, particularly the substantial declines for both Evo2 variants on pyrazinamide and the contrasting improvements for LogReg and SD-CNN on rifampicin.

#### I.3 Predictive Performance: Statistical Significance

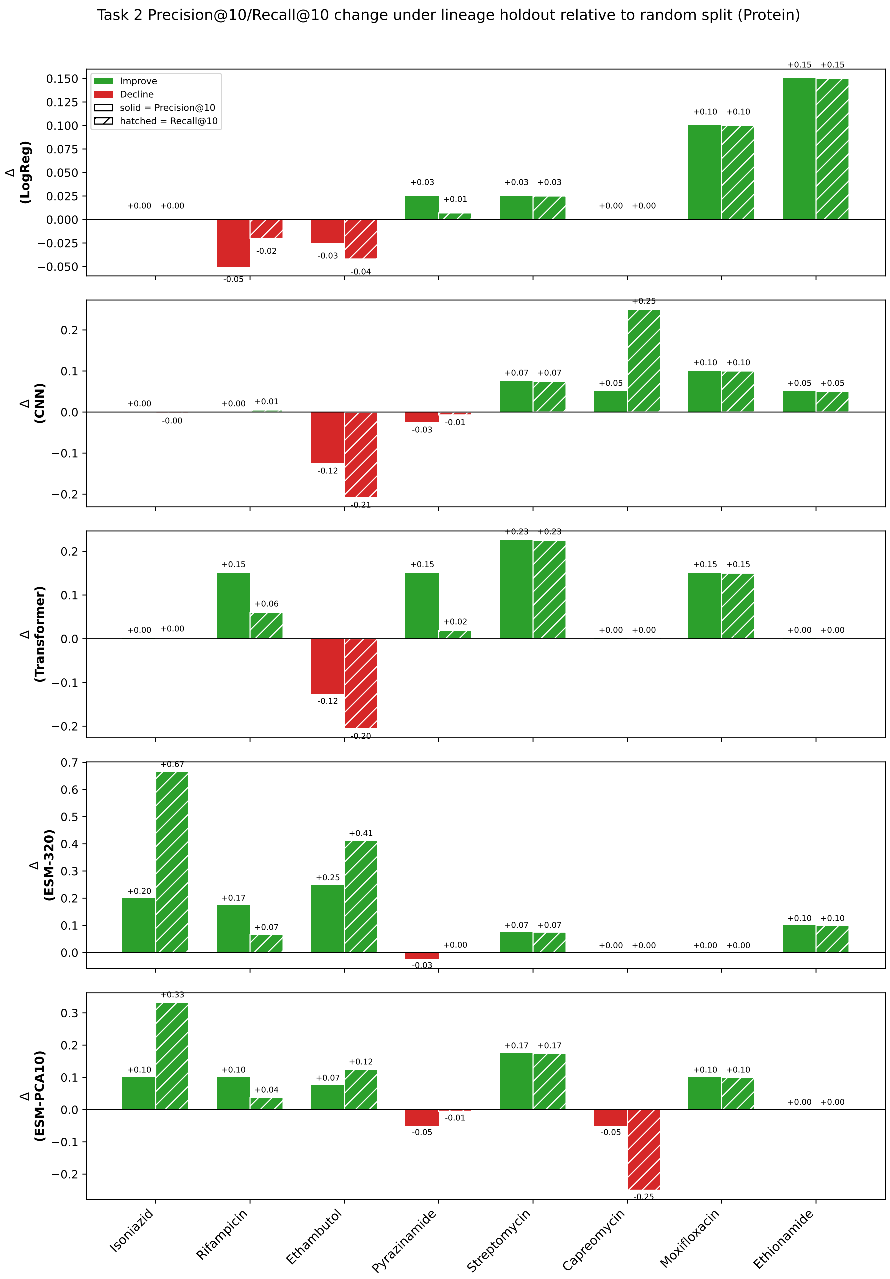


Figure 10: Per-model, per-drug change in Task 2 Precision@10 (left, solid bar) and Recall@10 (right, hatched bar) under lineage holdout relative to the random split, for the five protein models with a lineage-holdout value: LogReg, CNN, Transformer, ESM-320, ESM-PCA10. Amikacin (no protein-coding WHO target positions) and Kanamycin (non-coding locus) are excluded, matching Table 5.

Table 14: Isolate-level held-out train/test split used for the protein test AUCs in Table 2 (80/20, stratified by phenotype, seed 42). Each isolate appears in exactly one partition; for multi-gene drugs the split is performed after concatenating an isolate's genes, so no isolate contributes to both training and testing.

| | | Train | | | Test | | |
|---|---|---|---|---|---|---|---|
| Drug | Loci | $n$ | R | S | $n$ | R | S |
| Isoniazid | 2 | 13,948 | 4,660 | 9,288 | 3,488 | 1,165 | 2,323 |
| Rifampicin | 1 | 14,065 | 3,895 | 10,170 | 3,517 | 974 | 2,543 |
| Ethambutol | 3 | 12,088 | 2,356 | 9,732 | 3,023 | 589 | 2,434 |
| Pyrazinamide | 1 | 10,323 | 1,705 | 8,618 | 2,581 | 426 | 2,155 |
| Streptomycin | 2 | 6,145 | 1,933 | 4,212 | 1,537 | 483 | 1,054 |
| Amikacin | 1 | 2,964 | 566 | 2,398 | 741 | 141 | 600 |
| Capreomycin | 1 | 3,083 | 540 | 2,543 | 771 | 135 | 636 |
| Moxifloxacin | 2 | 2,294 | 310 | 1,984 | 574 | 78 | 496 |
| Levofloxacin | 2 | 215 | 61 | 154 | 54 | 15 | 39 |
| Ethionamide | 3 | 1,722 | 552 | 1,170 | 431 | 138 | 293 |

Table 15: Pooled comparison of DNA model performance against the Logistic Regression baseline, aggregated across drugs. For each drug and model, the five cross-validation fold AUCs were first averaged to a single per-drug mean AUC; a paired Wilcoxon signed-rank test (two-sided) was then applied across drugs ($n = 11$), pairing each model's per-drug mean AUCs against the corresponding per-drug Logistic Regression means. Holm–Bonferroni correction was applied across the four models in this table.

| Model vs. Baseline | Mean LogReg AUC | Mean Model AUC | ΔAUC | $p$ (Wilcoxon) | $p$ (Holm) | Num. Drugs |
|---|---|---|---|---|---|---|
| SD-CNN | 0.867 | 0.864 | -0.003 | 0.966 | 1.000 | 11 |
| SD-Evo2-CNN (frozen) | 0.867 | 0.871 | +0.004 | 0.765 | 1.000 | 11 |
| SD-Evo2-CNN-PCA10 (frozen) | 0.867 | 0.873 | +0.006 | 0.413 | 1.000 | 11 |
| MD-CNN | 0.867 | 0.922 | +0.055 | 0.001 | 0.004 | 11 |

Table 16: Pooled comparison of protein model performance against the Logistic Regression baseline, aggregated across drugs. For each drug and model, the five cross-validation fold AUCs were first averaged to a single per-drug mean AUC; a paired Wilcoxon signed-rank test (two-sided) was then applied across drugs ($n = 10$), pairing each model's per-drug mean AUCs against the corresponding per-drug Logistic Regression means. Holm–Bonferroni correction was applied across the four models in this table.

| Model vs. Baseline | Mean LogReg AUC | Mean Model AUC | ΔAUC | $p$ (Wilcoxon) | $p$ (Holm) | Num. Drugs |
|---|---|---|---|---|---|---|
| CNN | 0.799 | 0.785 | -0.014 | 0.037 | 0.074 | 10 |
| ESM-320 | 0.799 | 0.783 | -0.016 | 0.002 | 0.008 | 10 |
| ESM-PCA10 | 0.799 | 0.791 | -0.008 | 0.131 | 0.131 | 10 |
| Transformer | 0.799 | 0.630 | -0.169 | 0.002 | 0.008 | 10 |

Table 17: Significance test for the random-split-to-lineage-holdout AUC change reported in Table 4 (DNA). For each architecture, a paired Wilcoxon signed-rank test (two-sided) compared each drug's lineage-holdout mean AUC (averaged across eligible held-out lineages) against that drug's random-split test AUC (Table 2), pairing by drug across $n = 10$ drugs (excludes Levofloxacin, which did not satisfy the lineage-split criterion). Holm-Bonferroni correction was applied across the four architectures reported in Table 4.

| Architecture | Mean Random-Split AUC | Mean Lineage-Holdout AUC | ΔAUC | $p$ (Wilcoxon) | $p$ (Holm-corrected) |
|---|---|---|---|---|---|
| SD-LogReg | 0.8682 | 0.8322 | -0.0359 | 0.0488 | 0.1465 |
| SD-CNN | 0.8829 | 0.8347 | -0.0482 | 0.0645 | 0.1465 |
| Evo2-CNN (frozen) | 0.8700 | 0.8238 | -0.0462 | 0.0059 | 0.0234 |
| Evo2-CNN-pca10 (frozen) | 0.8625 | 0.8164 | -0.0461 | 0.0645 | 0.1465 |

Table 18: Significance test for the random-split-to-lineage-holdout AUC change reported in Table 4 (protein). For each architecture, a paired Wilcoxon signed-rank test (two-sided) compared each drug's lineage-holdout mean AUC (averaged across eligible held-out lineages) against that drug's random-split test AUC (Table 2), pairing by drug across $n = 9$ drugs (excludes Kanamycin, not protein-coding, and Levofloxacin, which did not satisfy the lineage-split criterion). Holm-Bonferroni correction was applied across the five architectures reported in Table 4.

| Architecture | Mean Random-Split AUC | Mean Lineage-Holdout AUC | ΔAUC | *p* (Wilcoxon) | *p* (Holm-corrected) |
|---|---|---|---|---|---|
| LogReg | 0.788 | 0.750 | -0.037 | 0.004 | 0.020 |
| CNN | 0.766 | 0.747 | -0.019 | 0.004 | 0.020 |
| Transformer | 0.614 | 0.602 | -0.012 | 0.734 | 0.734 |
| ESM-320 | 0.754 | 0.724 | -0.030 | 0.074 | 0.148 |
| ESM-PCA10 | 0.772 | 0.726 | -0.046 | 0.004 | 0.020 |

Table 19: Pooled significance test for Table 3 (DNA), comparing each model's WHO-catalogued resistance variant recovery against the Logistic Regression baseline. Table 3 reports single point-estimate Precision@10/Recall@10 values per drug, and here, a paired Wilcoxon signed-rank test (two-sided) was applied across drugs ($n = 11$), pairing each model's per-drug Precision@10 (and, separately, Recall@10) against Logistic Regression's. Holm-Bonferroni correction was applied within each metric's family of four models.

| | Precision@10 | | | | Recall@10 | | | |
|---|---|---|---|---|---|---|---|---|
| Model vs. LogReg | Mean LogReg | Mean Model | Δ | *p* (Holm) | Mean LogReg | Mean Model | Δ | *p* (Holm) |
| SD-CNN | 0.391 | 0.391 | +0.000 | 0.858 (0.858) | 0.206 | 0.250 | +0.045 | 0.260 (0.260) |
| MD-CNN | 0.391 | 0.136 | -0.255 | **0.005 (0.020)** | 0.206 | 0.076 | -0.129 | **0.005 (0.020)** |
| SD-Evo2-CNN (frozen) | 0.391 | 0.227 | -0.164 | 0.033 (0.066) | 0.206 | 0.122 | -0.084 | 0.086 (0.172) |
| SD-Evo2-CNN-PCA10 (frozen) | 0.391 | 0.200 | -0.191 | **0.008 (0.023)** | 0.206 | 0.095 | -0.110 | **0.008 (0.023)** |

Table 20: Pooled significance test for Table 3 (protein), comparing each model's WHO-catalogued resistance variant recovery against the Logistic Regression baseline. Table 3 reports single point-estimate Precision@10/Recall@10 values per drug, and here, a paired Wilcoxon signed-rank test (two-sided) was applied across drugs ($n = 10$), pairing each model's per-drug Precision@10 (and, separately, Recall@10) against Logistic Regression's. Holm-Bonferroni correction was applied within each metric's family of four models.

| | Precision@10 | | | | Recall@10 | | | |
|---|---|---|---|---|---|---|---|---|
| Model vs. LogReg | Mean LogReg | Mean Model | Δ | *p* (Holm) | Mean LogReg | Mean Model | Δ | *p* (Holm) |
| CNN | 0.420 | 0.490 | +0.070 | 0.175 (0.349) | 0.415 | 0.466 | +0.051 | 0.499 (0.749) |
| ESM-320 | 0.420 | 0.320 | -0.100 | 0.104 (0.312) | 0.415 | 0.203 | -0.212 | 0.080 (0.239) |
| ESM-PCA10 | 0.420 | 0.400 | -0.020 | 0.812 (0.812) | 0.415 | 0.358 | -0.057 | 0.374 (0.749) |
| Transformer | 0.420 | 0.260 | -0.160 | **0.012 (0.047)** | 0.415 | 0.203 | -0.212 | **0.012 (0.046)** |

Table 21: Significance test for the DNA Task 2 lineage-holdout ΔPrecision@10/ΔRecall@10 values reported in Table 5. For each architecture, a paired Wilcoxon signed-rank test (two-sided) compared each drug's lineage-holdout-controlled Precision@10 (and, separately, Recall@10) against that drug's random-split value from the same controlled-SHAP comparison, pairing by drug across $n = 10$ drugs. Holm-Bonferroni correction was applied within each metric's family of four architectures reported in Table 5.

| | Precision@10 | | Recall@10 | |
|---|---|---|---|---|
| Architecture | Mean Δ | *p* (Holm) | Mean Δ | *p* (Holm) |
| SD-LogReg | +0.0175 | 0.594 (1.000) | +0.0385 | 0.515 (1.000) |
| SD-CNN | +0.0300 | 0.512 (1.000) | -0.0219 | 0.275 (0.826) |
| SD-Evo2-CNN (frozen) | -0.1175 | 0.017 (0.069) | -0.0756 | 0.008 (0.031) |
| SD-Evo2-CNN-PCA10 (frozen) | -0.0375 | 1.000 (1.000) | +0.0139 | 0.674 (1.000) |

Table 22: Significance test for the protein Task 2 lineage-holdout ΔPrecision@10/ΔRecall@10 values reported in Table 5. For each architecture, a paired Wilcoxon signed-rank test (two-sided) compared each drug's lineage-holdout-controlled Precision@10 (and, separately, Recall@10) against that drug's random-split value from the same controlled-SHAP comparison, pairing by drug across $n = 8$ drugs. Holm-Bonferroni correction was applied within each metric's family of five architectures reported in Table 5.

| | Precision@10 | | Recall@10 | |
|---|---|---|---|---|
| Architecture | Mean Δ | *p* (Holm) | Mean Δ | *p* (Holm) |
| LogReg | +0.028 | 0.463 (0.924) | +0.028 | 0.345 (0.947) |
| CNN | +0.016 | 0.462 (0.924) | +0.033 | 0.383 (0.947) |
| Transformer | +0.069 | 0.074 (0.250) | +0.032 | 0.249 (0.947) |
| ESM-320 | +0.097 | 0.046 (0.232) | +0.166 | 0.028 (0.139) |
| ESM-PCA10 | +0.056 | 0.063 (0.250) | +0.065 | 0.237 (0.947) |

##### I.4 WHO-catalogued resistance Variant Recovery

Table 23: WHO variants recovered by the best CNN–OHE models at top-$k$=10. For each drug, the table lists the number of true positives (TP@10), the harmonic mean of precision and recall (F1@10), and representative recovered variants. DNA-based CNN results are shown for single-drug (SD) models, and protein-based CNN results are shown for per-drug models.

| DNA SD-CNN–OHE (Task 2:WHO-catalogued resistance Variant recovery) | | | |
|---|---|---|---|
| Drug | TP@10 | F1@10 | Representative Identified Variants |
| Isoniazid | 3 | 0.17 | *katG_p.Ser315Thr*, *inhA_c.-777C>T* |
| Rifampicin | 5 | 0.23 | *rpoB_p.Ser450Leu*, *rpoB_p.His445Asp* |
| Ethambutol | 7 | 0.54 | *embB_p.Met306Val*, *embB_p.Asp354Ala* |
| Pyrazinamide | 5 | 0.03 | *pncA_p.Asp49Gly* |
| Streptomycin | 4 | 0.07 | *rpsL_p.Lys43Arg*, *gid_p.Ala183Val* |
| Amikacin | 3 | 0.43 | *rrs_n.1401A>G* |
| Capreomycin | 2 | 0.09 | *rrs_n.1401A>G* |
| Kanamycin | 1 | 0.11 | *rrs_n.1401A>G* |
| Moxifloxacin | 5 | 0.30 | *gyrA_p.Asp94Ala* |
| Levofloxacin | 5 | 0.30 | *gyrA_p.Asp94Ala* |
| Ethionamide | 3 | 0.03 | *inhA_c.-77C>T* |
| Protein CNN–OHE (Task 2: WHO-catalogued resistance Variant) | | | |
| Isoniazid | 2 | 0.31 | *katG_p.Ser315Arg*, *inhA_p.Ser94Ala* |
| Rifampicin | 10 | 0.56 | *rpoB_p.Arg448Gln*, *rpoB_p.Asp435Ala* |
| Ethambutol | 6 | 0.75 | *embB_p.Asp328Tyr*, *embB_p.Asp354Ala* |
| Pyrazinamide | 10 | 0.19 | *pncA_p.Asp8Ala*, *pncA_p.Cys138Arg* |
| Streptomycin | 7 | 0.70 | *rpsL_p.Lys43Arg*, *gid_p.Lys88Arg* |
| Amikacin | 0 | 0.00 | – |
| Capreomycin | 1 | 0.17 | *tlyA_p.Gly232Asp* |
| Kanamycin | – | – | – |
| Moxifloxacin | 5 | 0.50 | *gyrA_p.Ala90Val*, *gyrA_p.Asp89Asn* |
| Levofloxacin | 7 | 0.70 | *gyrB_p.Asn499Asp*, *gyrB_p.Ser447Phe* |
| Ethionamide | 1 | 0.10 | *inhA_p.Ser94Ala* |

###### I.4.1 Per-variant support in Task 2 cohorts

Supplementary Data 1 file reports mutation-level support for the WHO Group 1 and 2 nucleotide/amino-acid substitutions represented in the protein Task 2 gene sets. Carrier counts were calculated from the model-ready isolate sequences for each drug, with resistant and susceptible carrier counts reported separately. Task 2 evaluates residue positions, so multiple WHO substitutions can map to a single evaluated position; the table therefore distinguishes position-level eligibility from observation of the exact nucleotide/amino-acid substitution. The machine-readable version, including carrier fractions and exact-variant observation flags, is provided as Supplementary Data 1.

#### J Use of large language models

Claude (Anthropic; Claude Code) was used during the revision of this manuscript to: (i) review and debug analysis code; and (ii) suggest edits to the manuscript text. All code changes were reviewed and run by the authors, and all text was reviewed and approved by the authors. The LLM was not used to generate data or results.
